# Overcoming brain-derived therapeutic resistance in HER2+ breast cancer brain metastasis

**DOI:** 10.1101/2024.02.19.581073

**Authors:** Danyyl Ippolitov, Yi-Han Lin, Jeremy Spence, Aleksandra Glogowska, Thatchawan Thanasupawat, Jason Beiko, Marc R. Del Bigio, Xin Xu, Amy Wang, Raul Calvo, Abhijeet Kapoor, Juan J Marugan, Mark J Henderson, Thomas Klonisch, Sabine Hombach-Klonisch

## Abstract

Brain metastasis of HER2+ breast cancer occurs in about 50% of all women with metastatic HER2+ breast cancer and confers poor prognosis for patients. Despite effective HER2-targeted treatments of peripheral HER2+ breast cancer with Trastuzumab +/-HER2 inhibitors, limited brain permeability renders these treatments inefficient for HER2+ breast cancer brain metastasis (BCBM). The scarcity of suitable patient-derived in-vivo models for HER2+ BCBM has compromised the study of molecular mechanisms that promote growth and therapeutic resistance in brain metastasis. We have generated and characterized new HER2+ BCBM cells (BCBM94) isolated from a patient HER2+ brain metastasis. Repeated hematogenic xenografting of BCBM94 consistently generated BCBM in mice. The clinically used receptor tyrosine kinase inhibitor (RTKi) Lapatinib blocked phosphorylation of all ErbB1-4 receptors and induced the intrinsic apoptosis pathway in BCBM94. Neuregulin-1 (NRG1), a ligand for ErbB3 and ErbB4 that is abundantly expressed in the brain, was able to rescue Lapatinib-induced apoptosis and clonogenic ability in BCBM94 and in HER2+ BT474. ErbB3 was essential to mediate the NRG1-induced survival pathway that involved PI3K-AKT signalling and the phosphorylation of BAD at serine 136 to prevent apoptosis. High throughput RTKi screening identified the brain penetrable Poziotinib as highly potent compound to reduce cell viability in HER2+ BCBM in the presence of NRG1. Successful in-vivo ablation of BCBM94- and BT474-derived HER2+ brain tumors was achieved upon two weeks of treatment with Poziotinib. MRI revealed BCBM remission upon poziotinib, but not with Lapatinib treatment. In conclusion, we have established a new patient-derived HER2+ BCBM in-vivo model and identified Poziotinib as highly efficacious RTKi with excellent brain penetrability that abrogated HER2+ BCBM brain tumors in our mouse models.

## INTRODUCTION

Brain metastasis is a fatal complication occurring in 50% of patients with HER2+ breast cancer (BC) and represents one of the most adverse scenarios of HER2+ BC progression (1, 2) as successful treatments are lacking (3). Targeted therapies approved for application in primary and metastatic HER2+ BC involve both monoclonal antibodies (Trastuzumab, Pertuzumab) and small molecule RTK inhibitors (RTKi) such as Lapatinib, Neratinib, and Tucatinib (3, 4). RTKis are generally recommended as a second or third-line regimen for advanced BC patients who were unresponsive or developed resistance to the anti-ErbB mAbs (4). Despite a higher blood-brain barrier penetrability, RTKi monotherapies lack clinical efficacy (4, 5). When used as a monotherapy, the reversible dual EGFR/ErbB2 RTKi Lapatinib showed only a marginal response rate of 2.6-6% in patients with brain metastasis (BM) of HER2+ BC (6, 7). Monotherapy with the covalent dual EGFR/ErbB2 RTKi Neratinib showed a similar (8%) poor efficacy (8), although both Lapatinib and Neratinib proved to be more effective when combined with the deoxycytidine derivative Capecitabine (8, 9) in patients with brain metastases of HER2+ BC.

ErbB1-4 engage in ligand-activated homo- and hetero-dimerization with 11 different EGF-like ligands (10). ErbB2 (HER2) overexpression drives malignant transformation in BC as the preferred dimerization partner (11) for other RTKs of the ErbB family to activate downstream signaling via several pathways (e.g., PI3K/Akt, MAPK/ERK, PLCγ) known to mediate cell survival, proliferation, epithelial-mesenchymal transition, cell migration and tissue invasion (12). ErbB2-ErbB3 heterodimerization provides the most significant signaling in HER2+ BC (13, 14).

Neuregulins (NRG) constitute the largest subgroup of structurally related ErbB ligands (15-17). NRG1 specifically binds to the extracellular domain of ErbB3 and ErbB4 which results in formation of potentially six active ErbB dimers (12, 18). The role of NRGs in cancer progression is tightly linked to the ErbB-driven signaling in breast cancer (19-21). NRGs are commonly expressed in the central nervous system (22). NRG1 protein is expressed mainly by neurons, but also astrocytes, oligodendrocytes and microglia which are main resident cell types of the brain (https://www.proteinatlas.org/) (23). The expression of ADAM sheddases by brain resident cell populations may contribute to the release and paracrine HER2 activation by NRGs resulting in the progression of brain metastatic HER2+ BC at the brain metastatic niche (24, 25). Studies on breast-to-brain metastasis (BCBM) are hampered by the few patient-derived HER2+ BC cell models capable of generating brain metastases in mice (26), which severely limits mechanistic and therapeutic studies on BCBM.

The small molecule RTKi Lapatinib induces apoptosis in several HER2+ BC cell models (27) but several mechanisms of resistance to RTKis have been identified (28), among them the activation of compensatory pathways involving transcriptional and posttranslational up-regulation of ErbB3 (29), re activation of Akt (30), and PI3K independent induction of mTOR activity (31). For brain metastatic lesions, the expression of NRG1 in the brain (22, 32) can potentially mediate any of these mechanisms by binding to ErbB3/4 and inducing ErbB activation. ErbB3-PI3K-Akt signaling is known to mediate the anti-apoptotic response through the regulation of Bcl-2 and IAP protein families (33-35). Although endogenous NRG1 was shown to mediate resistance of HER2+ BC to both Lapatinib (27) and Trastuzumab (36) through the activation of ErbB3 and concomitant downregulation of apoptosis, a mechanistic explanation of the anti-apoptotic response driven by NRG1 is still lacking. Here we introduce a novel patient-derived hematogenic HER2+ BC brain metastasis model (BCBM94) and identify NRG1-driven mechanisms that rescue Lapatinib-induced apoptosis. We show that NRG1 fails to rescue apoptosis by the brain penetrable and irreversible ErbB1/2/4 RTKi Poziotinib and demonstrate the ability of Poziotinib to abrogate with high efficacy HER2+ BC metastatic brain lesions in mice.

## RESULTS

### Establishment and characterization of a new BCBM94 cell and mouse model

A neurosurgical tissue sample was obtained from a metastatic adenocarcinoma to the cerebellum of a female patient diagnosed with invasive ductal carcinoma (grade III, stage T2N1). This research was approved by the Health Research Ethics Board (HREB; protocol #19-038), University of Manitoba. The original patient breast tumor was identified as luminal-B HER2+ BC expressing HER2+ (score 3+), ER+ (Allred score 7), and devoid of PRα/β. Ultrasound-guided xenografting of BCBM94 cells into the left ventricle of Rag2γc-/- mice robustly led to brain metastasis in mice about 3 months after xenografting. The ability of BCBM94 cells to establish hematogenic brain metastases was confirmed in three consecutive rounds of isolation of tumor cells from mouse brain and intracardial re-injection for brain colonization. BC metastatic lesions were of epithelial morphology and formed vascularized and proliferative metastases of different sizes throughout the brain as determined by immunoreactive CD31 and Ki67, respectively (**Fig. 1A)**. Strong membrane expression of ErbB2 was detected in the patient primary BC tissue and corresponding brain metastatic tissue, as well as in brain metastases of the experimental animals **(Fig. 1B).** Membrane expression of ErbB2 was also detected in cultured BCBM94 cells **(Fig. 1C)**. When compared to cultured triple-negative MDA-MB-BR or HER2+ cell lines BT474 and SKBR3, BCBM94 showed the highest expression of ErbB2 protein, low level of ERα, and complete lack of PRα/β proteins **(Fig. 1D)**. These findings identified BCBM94 cells as novel luminal-B HER2+ BC subtype capable of hematogenous brain metastasis **(Fig. 1D)**.

**Figure 1.**
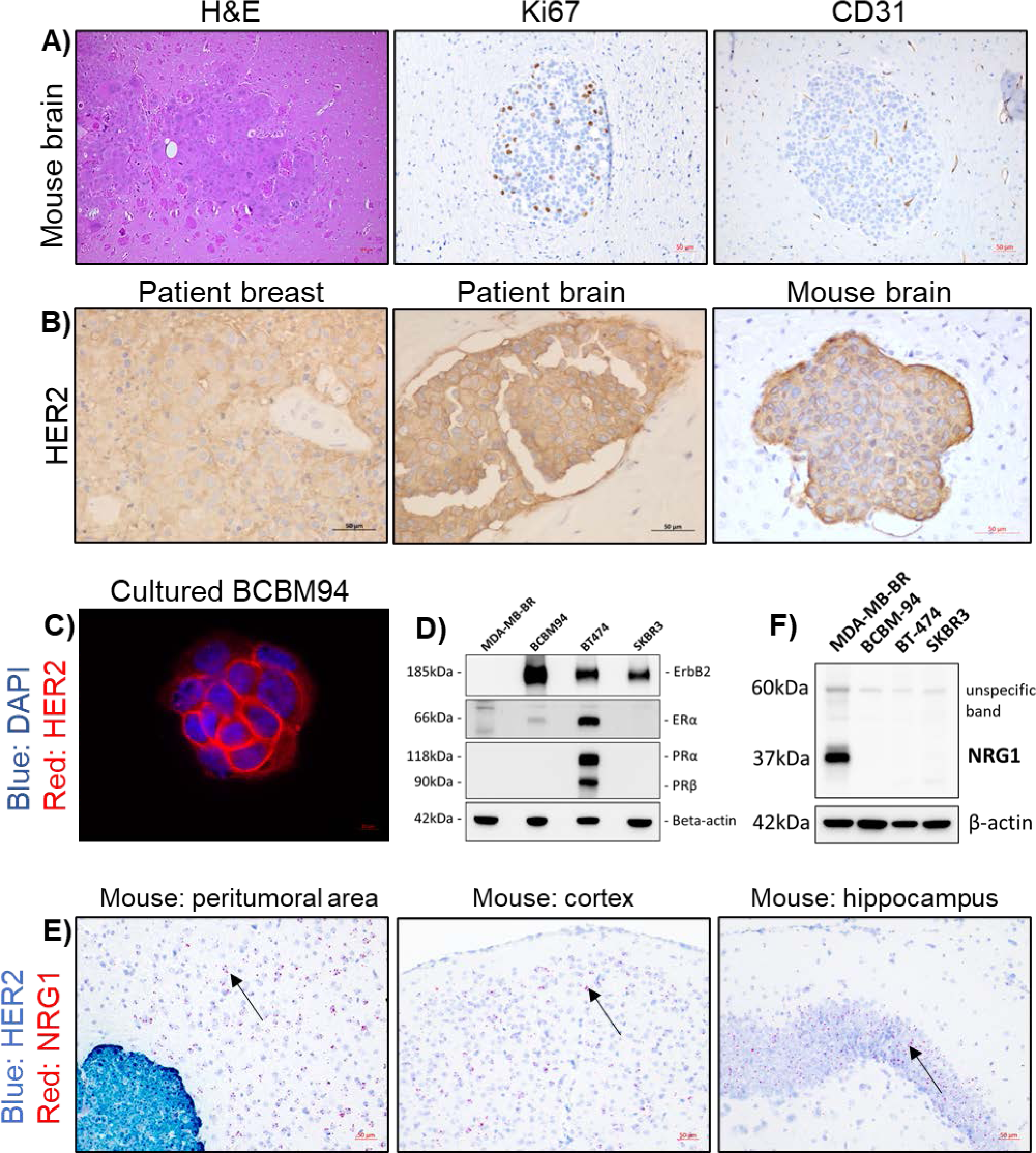
HER2+ BCBM94 cells establish hematogenic brain metastasis in mice. **(A)** BCBM94 cells produce hematogenous brain metastasis upon intracardiac xenografting in RAG2yc-/- mice. H&E and IHC staining for Ki67+ nuclei and CD31+ endothelial cells in FFPE mouse brain tissues containing BCBM94 lesions. Magnification 200x (**B,C)** BCBM94 cells retain HER2 expression *in-vivo* and *in-vitro.* HER2 IHC staining of FFPE tissue sections of patient’s breast, patient’s brain, and mouse brain containing BCBM94 lesions. Magnification 200x. ICC staining of cultured formaldehyde fixed BCBM94 cells show strong membrane expression of HER2 (ErbB2). (**D)** BCBM94 is a Luminal B HER2+ BC model. Western blot (WB) comparing total protein expression of HER2 (ErbB2), ERα, and PRα/β in the triple-negative MDA-MB-BR and HER2+ BCBM94, BT474, and SKBR3 cell lines *in vitro. (***E)** NRG1 is expressed in the TME of BCBM94 brain metastases. *In-situ* expression of NRG1 and ErbB2 mRNA in the FFPE mouse brain tissue containing BCBM94 metastases was assessed with the RNAscope 2.5 HD Duplex assay. Black arrows indicate NRG1 mRNA (red dots) within non-tumoral cells in the mouse brain. NRG1+ cells are abundantly present in various regions of the mouse brain, including the TME of HER2+ BCBM94 (blue) metastasis. Magnification 200x. (**F)** HER2+ BC cell models are devoid of NRG1. WB compared NRG1 protein expression in the triple-negative MDA-MB BR and HER2+ BCBM94, BT474, and SKBR3 cell lines *in-vitro*.

*In-situ* hybridization of BCBM94 metastatic mouse brain revealed abundant mouse *Nrg1* mRNA expression in different regions, including mouse brain tissue adjacent to BCBM94 brain metastatic lesions, which were highly positive for human ErbB2 transcripts **(Fig. 1E)**. Western blot analysis showed the absence of endogenous NRG1 protein expression in protein lysates of BCBM94 and two other HER2+ BC cell lines (BT474, SKBR3), whereas triple-negative MDA-MB231BR cells had detectable level of NRG1 protein **(Fig. 1F)**. Quantitative RT-PCR detected transcripts for ErbB1-4 but weak to undetectable levels of NRG1-4 transcripts in BCBM94 cells **(Sup. Fig. 1A)**. These results suggested ErbB3/4+ BCBM94 as a target of Nrg1 produced by the mouse brain.

### NRG1 counteracts Lapatinib cytotoxicity in HER2+ brain metastatic BC cells

We evaluated the effects of the small molecule EGFR/ErbB2 RTKi Lapatinib and recombinant human NRG1 **(rhNRG1)** on the viability of HER2+ brain metastatic BC cells BCBM94 and BT474. WST-1 cell viability assays showed a dose-dependent decline in BCBM94 cell numbers upon treatment with Lapatinib. The half maximal inhibitory concentration **(IC_50_)** of Lapatinib in WST-1 assays for BCBM94 cells was reached at 250nM **(Fig. 2A)**. At this IC_50_ Lapatinib concentration, increasing concentrations of rhNRG1 resulted in a dose-dependent rescue of cell viability. This rescue effect was maximal at 5ng/mL rhNRG1 **(Fig. 2B)**, which is an NRG1 concentration reported in human serum (37). From hereon, 5ng/mL rhNRG1 was used for all *in vitro* experiments. When combined with Lapatinib at IC_50_, rhNRG1 maintained the viability of BCBM94 cells at 80-90% of control values as determined at 24-72h by WST assays **(Fig. 2C)**.

**Figure 2.**
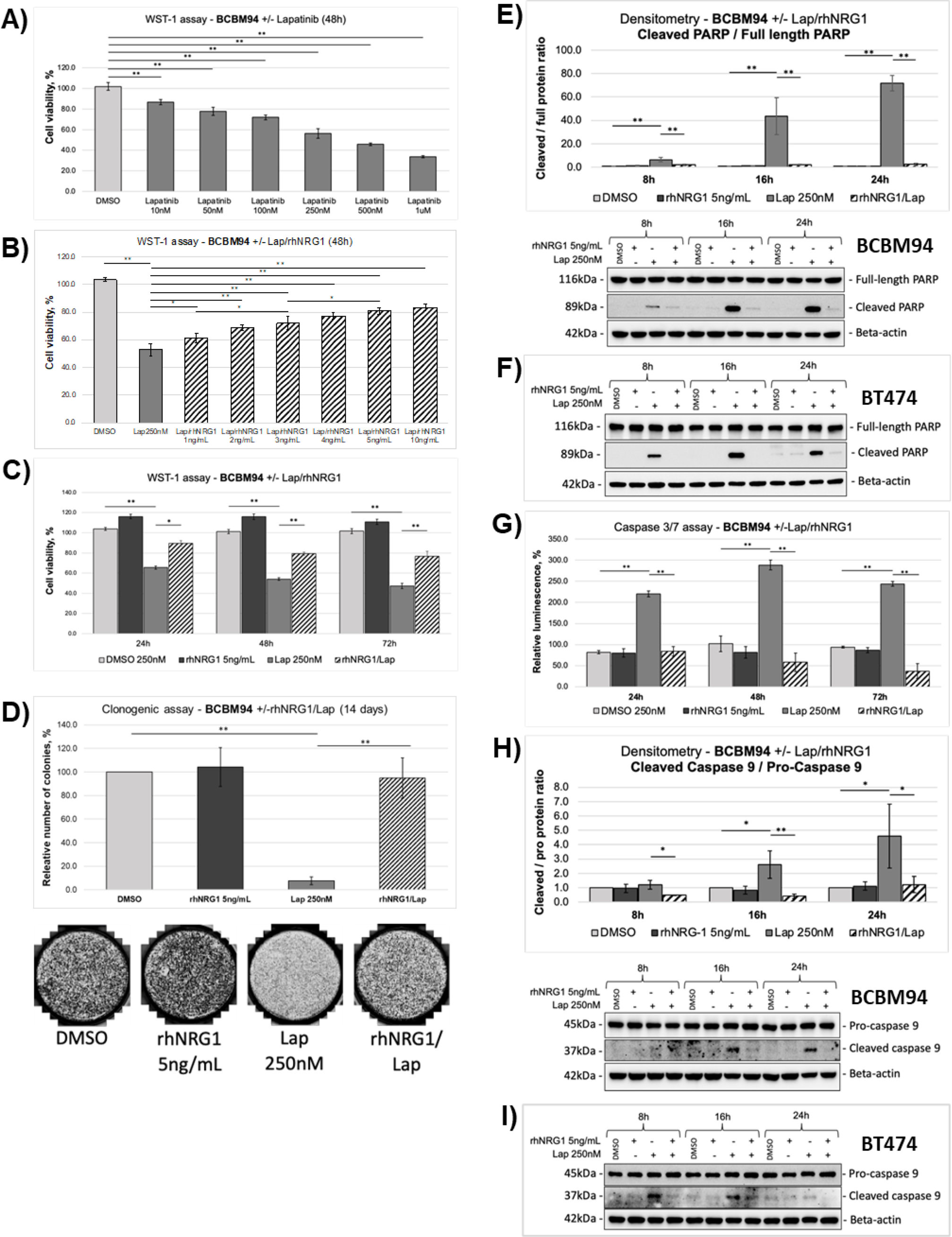
NRG1 rescues BCBM94 cells from Lapatinib-induced cytotoxicity. **(A-D)** Cell viability of BCBM94 under Lapatinib (Lap) +/- rhNRG1 treatment was assessed in the WST 1 assay. The endpoint absorbance readouts were used for quantification of the relative cell viability (mean +/- SD, n=3) **(A-C).** Cell proliferation and colony formation potential of BCBM94 cells under Lap +/- rhNRG1 treatment were assessed in colony formation assays **(D).** The bar chart presents the average number of colonies formed by BCBM94 cells under Lap +/- rhNRG1 treatment (n=2) **(D)**. Representative images of colonies are shown below the plot **(D)**. **(E-I)** NRG1 counteracts Lapatinib-induced apoptosis. The chart presents cleaved / full length PARP protein ratio in BCBM94 cells under Lap +/-rhNRG1 conditions measured by Western blot (WB) and quantified with densitometry (n=3) **(E)**. A representative WB is shown below the plot **(E).** WB detection of cleaved and full-length PARP proteins in BT474 cells is shown under Lap +/- rhNRG1 conditions (n=1) **(F)**. Relative luminescence values represent the activity of caspase-3 and caspase-7 under Lap +/-rhNRG1 conditions measured using a CaspaseGlo 3/7 assay (mean +/- SD, n=3) **(G).** Detection of the cleaved/ pro- caspase-9 protein ratio in BCBM94 cells under Lap +/-rhNRG1 conditions was measured by WB and quantified with densitometry (n=3) **(H)**. A representative WB is shown below the chart **(H).** WB detection of cleaved / pro- caspase-9 proteins in BT474 cells under Lap +/- rhNRG1 conditions (n=1) **(I).** Bar charts present mean +/- SD,*p<0.05, **p<0.01.

Measuring cellular impedance as a function of cell number and proliferation rate, xCELLigence Real Time Cell Analysis assays confirmed the ability of rhNRG1 to rescue BCBM94 from Lapatinib-induced cytotoxicity **(Sup. Fig. 1B, C)**. These findings were further validated in BT474 cells where the cytoprotective action of rhNRG1 was even more pronounced **(Sup. Fig. 1D)**.

To evaluate the long-term effects of Lapatinib +/-rhNRG1 treatment on the proliferation of BCBM94, we performed colony formation assays. While treatment with 250 nM Lapatinib for 14 days reduced the number of cell colonies by 90% from colony numbers in the control, combined Lapatinib/ rhNRG1 treatment resulted in colonies numbers similar to the control **(Fig. 2D)**. NRG1 alone did not significantly alter colony numbers **(Fig. 2D)**. Western blot analysis revealed that the rescue effect was mediated exclusively by exogenous rhNRG1 as Lapatinib did not induce the upregulation of endogenous NRG1 transcripts or protein in BCBM94 cells **(Sup. Fig. 1E)**.

### NRG1 abolishes Lapatinib cytotoxicity by attenuating apoptosis

Next, we asked whether the observed viability changes involved regulation of apoptosis. BCBM94 and BT474 HER2+ BC cell models responded to Lapatinib with an increase in cleavage of the active catalytic domain of PARP, but cleaved PARP levels remained undetectable with combined Lapatinib/ rhNRG1 treatment **(Fig. 2E, F)**. Notably, the sequential addition of rhNRG1 secondary to exposure to Lapatinib for 24h was able to mitigate a Lapatinib-induced cleaved PARP induction **(Sup. Fig. 2A, B)**. To explore the events preceding the cleavage of PARP, we evaluated the activity of the effector caspase-3 and caspase-7. Caspase-Glo 3/7 assays showed a prominent activation of caspase-3/7 in BCBM94 cells exposed to Lapatinib. The observed induction of these pro-apoptotic caspases of the intrinsic apoptosis pathway was fully abrogated by rhNRG1 at 5 ng/ml **(Fig. 2G)**. Further investigation of the upstream steps of the apoptotic cascade revealed that Lapatinib mediated activation of the apoptosis-initiating caspase-9 in BCBM94 and BT474 cells which was completely abolished by rhNRG1 **(Fig. 2H, I)**. These results identified mitochondrial apoptotic pathways as a target of the anti-apoptotic action of rhNRG1.

### Anti-apoptotic action of NRG1 is mediated through the mitochondrial pathway

Members of the Bcl-2 family mainly regulate the intrinsic apoptotic pathway (38). In BCBM94 and BT474 cells, rhNRG1 rescued the Lapatinib-induced de-phosphorylation of the BH3-only protein Bad at Ser136, which is an Akt phosphorylation site **(Fig. 3A, B)**. The ability of the BH3-only and pore forming proteins of the Bcl-2 family to trigger apoptosis is determined not solely by their expression levels but also by cellular localization, phosphorylation, and dimerization status of these proteins (39). While we did not observe significant changes in the total levels of the Bcl-2 proteins **(Suppl. Fig. 2C F)**, cellular immunofluorescence analysis identified the appearance of Bax aggregates that co-localized with mitochondrial Bak in BCBM94 cells exposed to Lapatinib **(Fig. 3D)**. RhNRG1 preserved the diffuse cellular distribution of Bax and blocked formation of pro-apoptotic Bax/ Bak dimers observed in the presence of Lapatinib only **(Fig. 3D)**. MitoTracker assays showed a reduction of fluorescence signal intensity in Lapatinib-treated BCBM94 indicative of a lower number of active mitochondria **(Fig. 3C)** and rhNRG1 was able to mitigate this Lapatinib effect on mitochondria **(Fig. 3C)**. TEM ultrastructural imaging revealed prominent damage to mitochondrial cristae and ruptured outer mitochondrial membranes in BCBM94 cells treated with Lapatinib **(Suppl. Fig. 2G)**, suggestive of mitochondrial damage. Like in control cells, these morphological alterations were only rarely observed in cells receiving dual Lapatinib/ rhNRG1 treatment **(Suppl. Fig. 2G)**. We concluded that rhNRG1 rescued apoptosis by blocking the pro-apoptotic cascade at the outer mitochondrial membrane.

**Figure 3.**
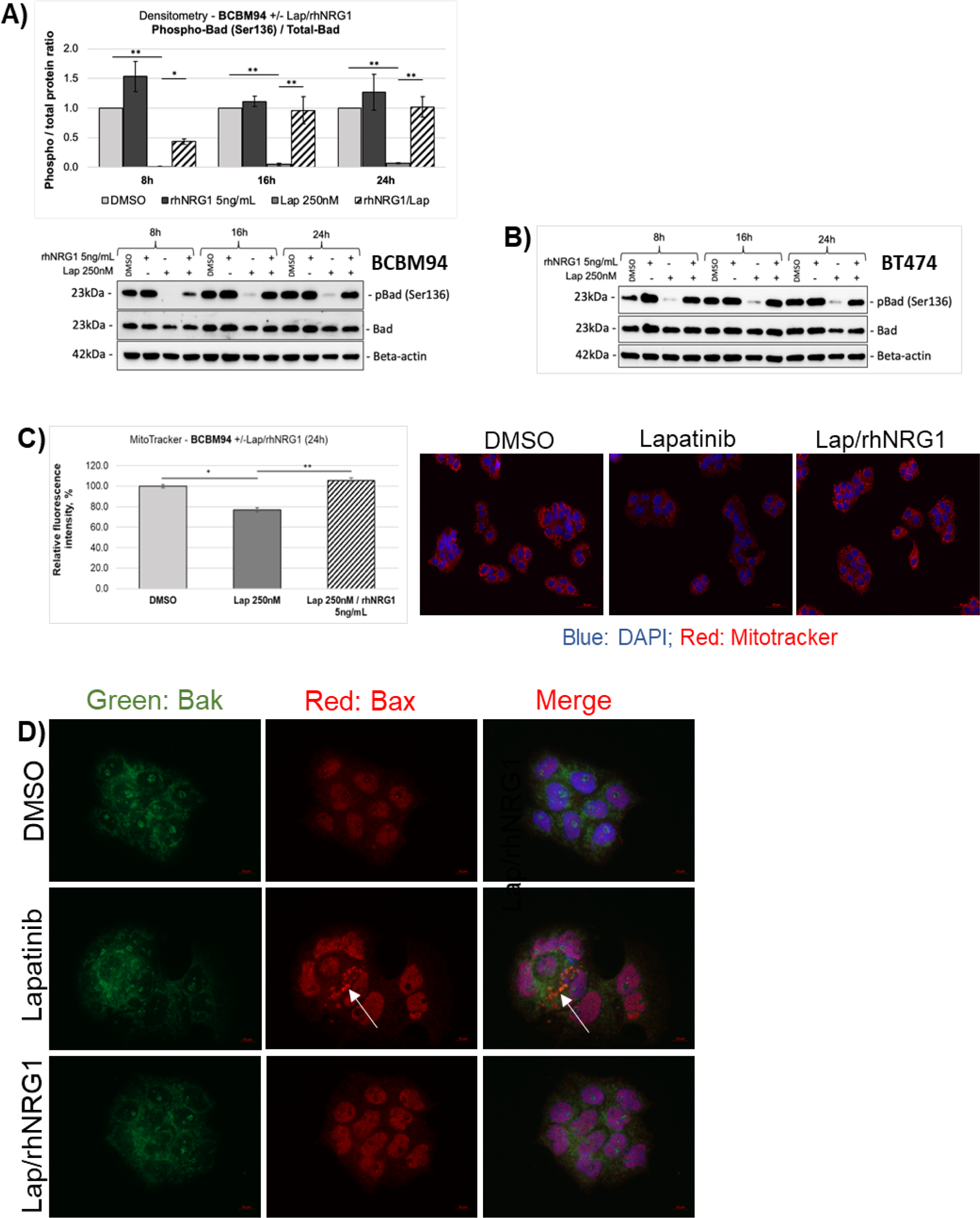
Anti-apoptotic actions of NRG1 involve BCL2 proteins. **(A,B)** NRG1 rescues Bad phosphorylation under Lapatinib (Lap). The phospho-/ total Bad protein ratio in BCBM94 cells under Lap +/-rhNRG1 conditions was determined by WB and quantified with densitometry (n=3) **(A).** A representative WB is shown below the chart **(A).** WB images show expression of phospho- and total Bad proteins in BT474 cells under Lap +/- rhNRG1 conditions (n=1) **(B). (C)** NRG1 protects mitochondria from Lapatinib-induced damage. Relative fluorescence intensity values represent the number of active mitochondria in BCBM94 cells under Lap +/-rhNRG1 conditions detected with MitoTracker^®^ (n=2). Representative IF images are shown below the graph, magnification 200x **(C)**. (**D)** NRG1 prevents aggregation of the mitochondria outer membrane pore-formers Bax and Bak under Lapatinib. BCBM94 cells treated with Lap +/-rhNRG1 were PFA-fixed for detection of Bak and Bax by ICC/IF. The white arrow indicates punctate Bax aggregates co-localizing with Bak under Lapatinib treatment (n=2). Magnification 630x. Bar charts present mean +/- SD, *p<0.05, **p<0.01.

### NRG1 utilizes an ErbB3-Akt pathway to promote Lapatinib resistance

In both BCBM94 and BT474 cell models, Lapatinib potently decreased phosphorylation of ErbB1 to 4 and reduced total ErbB1 and ErbB4 protein content **(Fig. 4A-D)**. rhNRG1 exclusively rescued ErbB3 phosphorylation in both cell lines **(Fig. 4A-D)**. Intriguingly, the patient tissues derived from the primary breast tumor and the brain metastatic tissues used to isolate BCBM94 cells contained immunoreactive phosphorylated ErbB3, as did BCBM94 metastatic lesions in mouse brain **(Fig. 4E).** The presence of constitutively activated ErbB3 *in situ* and the ability of rhNRG1 to rescue ErbB3 phosphorylation upon Lapatinib treatment suggested a key role of ErbB3 in NRG1-mediated survival of both HER2+ BC models. While selective siRNA-mediated ErbB3 knockdown (KD) alone did not induce apoptosis, treatment of BCBM94**^ErbB3-KD^** and BT474**^ErbB3-KD^** cells with Lapatinib resulted in a significantly higher level of PAPR cleavage compared to mock transfected cells treated with Lapatinib **(Fig. 5A, B)**. ErbB3 KD also attenuated the ability of rhNRG1 to counteract Lapatinib-induced PARP cleavage in BCBM94 and BT474 cells **(Fig. 5A, B)**. ErbB3 KD also markedly weakened the ability of rhNRG1 to rescue BAD^Ser136^ phosphorylation under Lapatinib. Significantly reduced cellular levels of phospho-BAD^Ser136^ were detected upon co-treatment with Lapatinib and rhNRG1 compared to mock transfected BCBM94/ BT474 cells **(Fig. 5C, D)**. Lapatinib further decreased BAD^Ser136^ phosphorylation in both ErbB3-KD cell models. While BCBM94^ErbB3-KD^ and BT474^ErbB3-KD^ cells showed a small increase in phospho-BAD^Ser136^ upon NRG1, overall phospho-BAD levels were negligible compared to mock silenced BCBM94 and BT474 cells co-treated with Lapatinib and rhNRG1 **(Fig. 5C, D)**. Phosphorylation of BAD at Ser136 is mediated by activated AKT (33) and the rhNRG1-mediated rescue of ErbB3 phosphorylation under Lapatinib coincided with increased phosphorylation of the ErbB downstream effector kinase AKT in BCBM94 and BT474 cells **(Fig. 5E, F)**. To demonstrate a causal involvement of AKT in the anti- apoptotic regulation by NRG1, we utilized the PI3K/AKT inhibitor PI-103. PI-103 abrogated Akt phosphorylation with a moderate downregulation of total AKT protein in untreated BCBM94 and BT474 cells and completely blocked the pAKT rescue by rhNRG1 in Lapatinib exposed cells **(Fig. 5G, H).** PI 103 alone increased PARP cleavage in BCBM94 and BT474 cells and this coincided with a complete loss of BAD^Ser136^ phosphorylation and an inability of NRG1 to rescue cells from apoptotic actions upon co-treatment with Lapatinib/ rhNRG1 **(Fig. 5G, H)**. Hence, in our HER2+ BCBM94 and BT474 models NRG1 utilizes an ErbB3-PI3K-AKT-BAD signaling cascade to cause resistance to Lapatinib (**Fig.6**).

**Figure 4.**
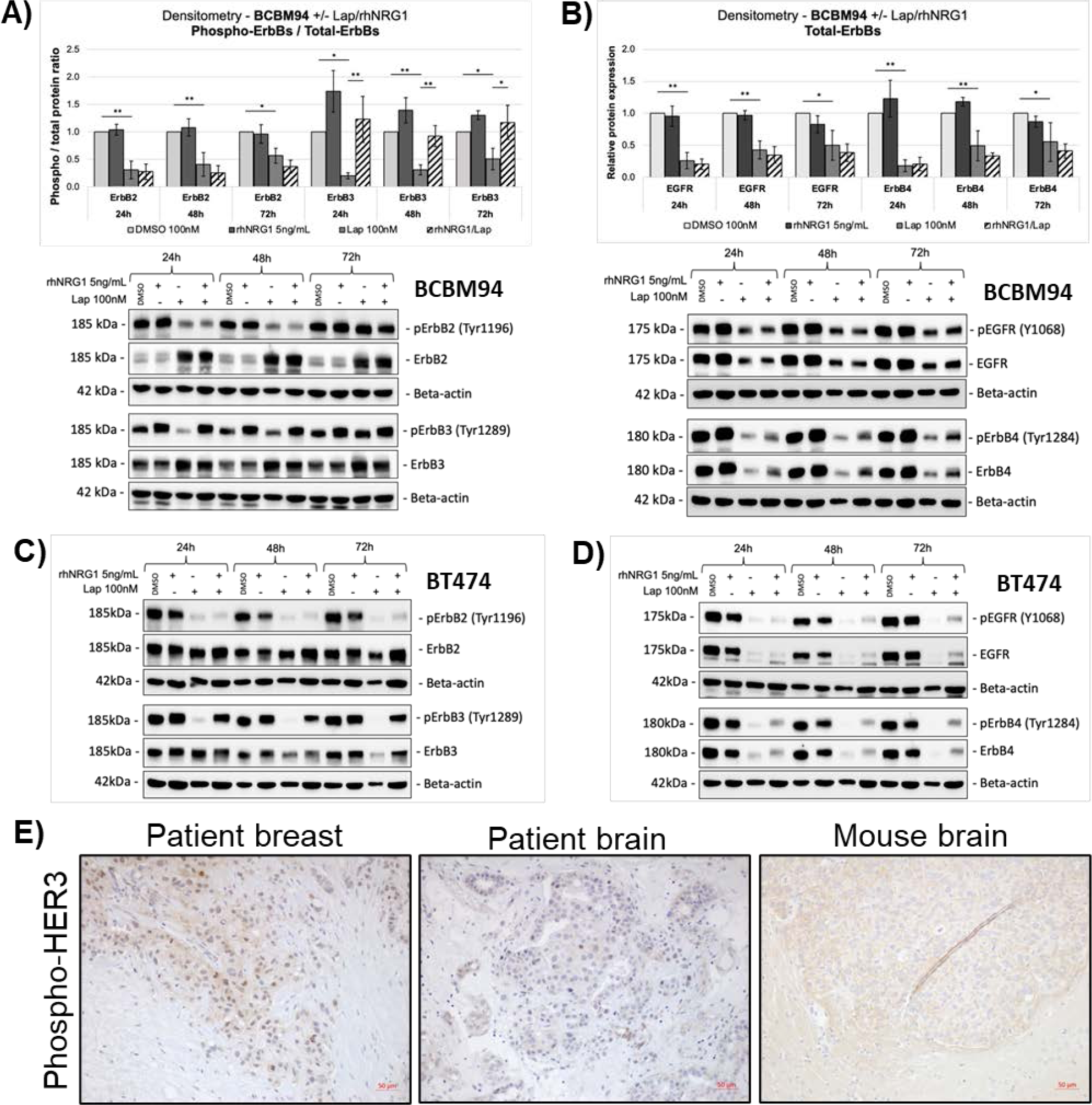
NRG1 rescues ErbB3 phosphorylation under HER2 inhibition in BCBM cells. **(A-D)** NRG1 rescues ErbB3 phosphorylation under Lapatinib (Lap). Protein ratios for phospho-/ total ErbB2 and phospho-/ total ErbB3 were determined in BCBM94 cells under Lap +/-rhNRG1 by WB and quantified with densitometry (n=3) **(A).** Total ErbB1 (EGFR) and total ErbB3 protein levels in BCBM94 cells under Lap +/-rhNRG1 conditions were measured by WB and quantified with densitometry (n=3) **(B).** Representative WBs are shown below the charts **(A, B).** WB images show the detection of phospho-ErbB1-4 and total ErbB1-4 proteins in BT474 cells under Lap +/-rhNRG1 conditions (n=1) **(C,D)**. (**E)** Phosphorylated ErbB3 is expressed by BCBM94 tumors *in-vivo*. IHC analysis of patient’s breast, patient’s brain, and mouse brain FFPE tissue sections containing BCBM94 lesions, magnification 200x. Bar charts present mean +/- SD, *p<0.05, **p<0.01.

**Figure 5.**
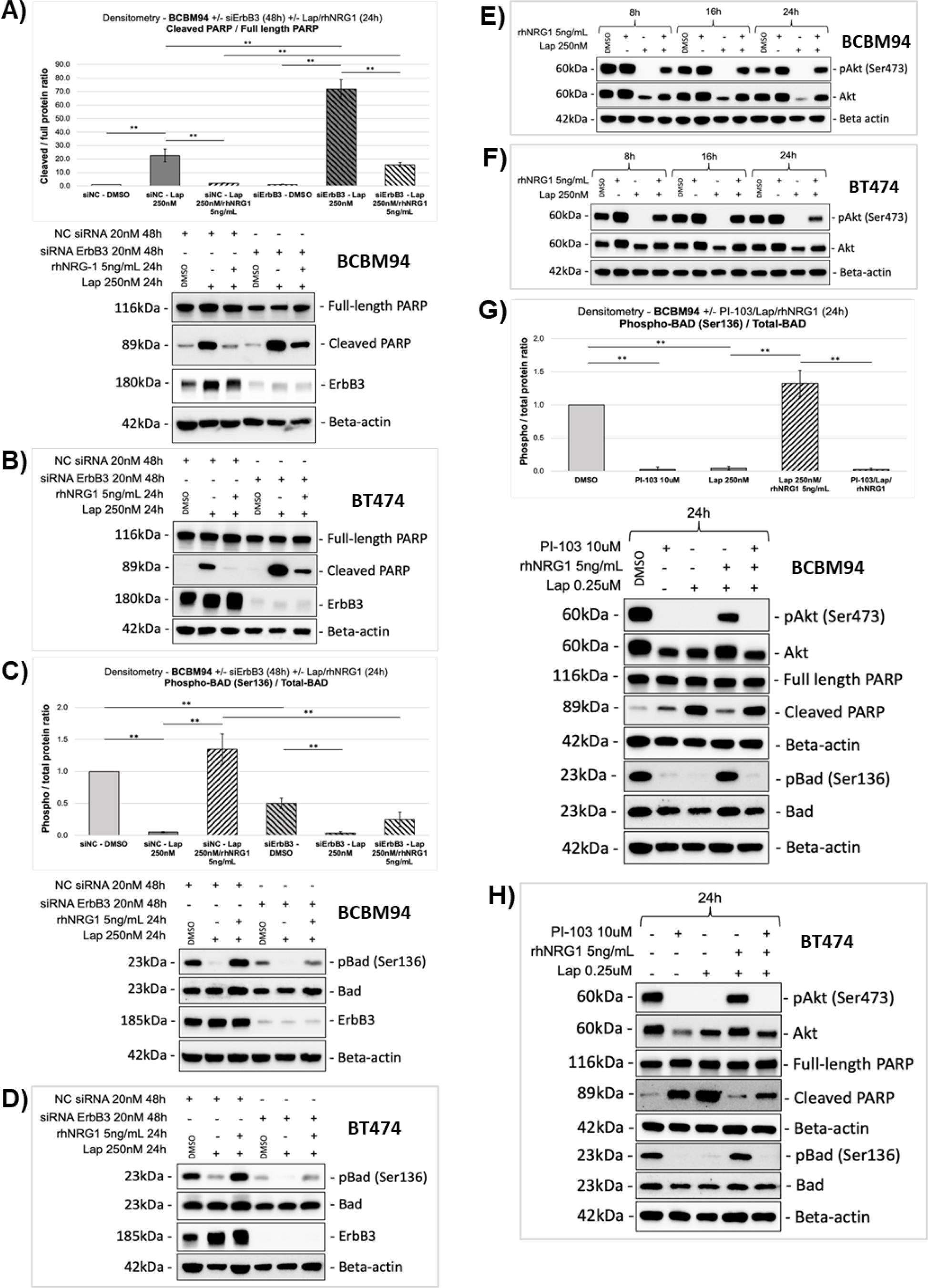
ErbB3 signaling mediates anti-apoptotic actions of NRG1 under Lapatinib. **(A, B)** Knockdown of ErbB3 increases PARP cleavage under Lapatinib (Lap) and mitigates NRG1 rescue. The cleaved/ full-length PARP protein ratio in BCBM94 cells under Lap +/-rhNRG1 +/-ErbB3siRNA conditions was measured by WB and quantified with densitometry (n=3) **(A).** A representative WB is shown below the chart **(A)**. Protein levels of cleaved and full-length PARP proteins in BT474 cells under Lap +/-rhNRG1 +/-ErbB3siRNA conditions are shown by WB (n=1) **(B). (C, D)** Knockdown of ErbB3 attenuates rhNRG1-mediated rescue of phospho-Bad under combined Lap/ rhNRG1 treatment. The graph presents phospho-/ total Bad protein ratio in BCBM94 cells under Lap +/-rhNRG1 +/- ErbB3siRNA conditions measured by WB and quantified with densitometry (n=3) **(C).** A representative WB is shown below the chart **(C)**. Protein levels of phospho- and total Bad proteins in BT474 cells under Lap +/-rhNRG1 +/-ErbB3siRNA conditions were detected by WB (n=1) **(D)**. (**E, F)** NRG1 rescues expression and phosphorylation of Akt under Lapatinib. WB images show expression of phospho- and total Akt proteins in BCBM94 (representative examples, n=3) **(E)** and BT474 (n=1) **(F)** cells under Lap +/-rhNRG1 +/-ErbB3siRNA conditions. (**G, H)** The anti-apoptotic action of NRG1 is mediated through Akt. The graph presents phospho-/ total Bad protein ratio in BCBM94 cells under Lap +/-rhNRG1 treatment measured by WB and quantified with densitometry; the PI3K inhibitor PI-103 was used at 10 µM **(G).** Representative WB images are shown below the chart (n=3) **(G)** and present the levels of phospho-/ total-Akt and cleaved/ full-length PARP proteins in BCBM94. Protein levels of phospho-/ total-Akt, cleaved/ full-length PARP and phospho-/ total Bad proteins in BT474 cells under Lap +/- rhNRG1 exposure and treatment with PI-103 were determined by WB (n=1). Bar charts present mean +/- SD, **p<0.01

**Figure 6.**
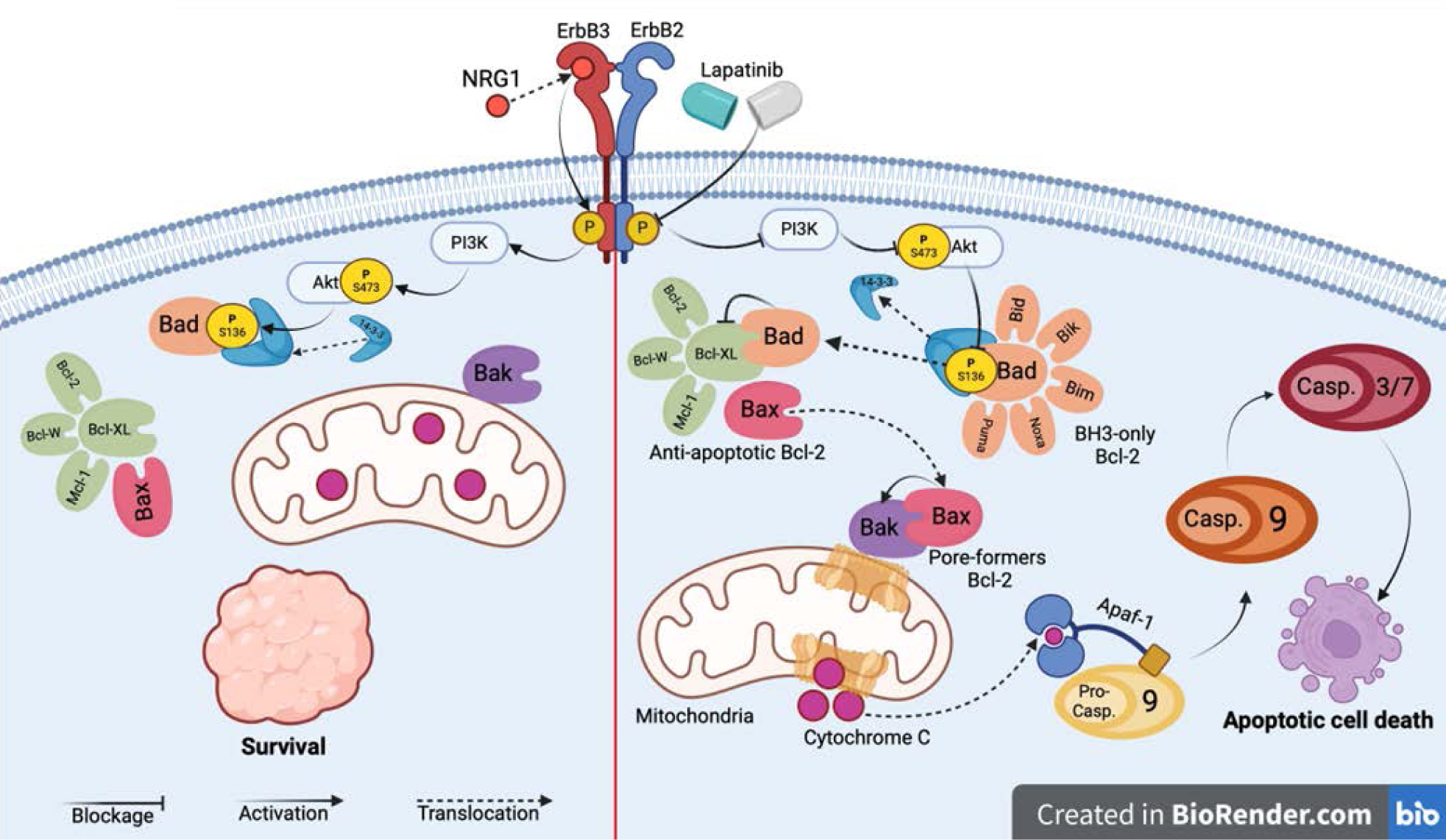
Schematic illustration of the NRG1 actions that rescue Lapatinib-induced apoptosis in HER2+ BCBM cells.

### NRG1 fails to rescue Poziotinib cytotoxicity in HER2+ BC *in-vitro*

In search of small molecule ErbB inhibitors capable of overcoming NRG1 RTKi resistance, we assembled a collection of 50 ErbB inhibitors, covering a range of isoform selectivities, mode of action (reversible vs covalent), and predicted CNS penetrance (**Suppl. Table 1**). We then tested the entire collection of compounds at 22-concentrations, ranging from 0.003 pM to 40 uM in the BCBM94 and HME1 immortalized human normal mammary epithelial cells using a CellTiterGlo viability assay. The dose-response profiles were compared to identify drug candidates that showed high selectivity and efficacy towards BCBM94, but not HME1 cells and were insensitive to the anti-apoptotic rescue by rhNRG1. This screen included the FDA-approved Lapatinib, Neratinib, and Tucatinib currently used as second line treatments in brain metastatic BC patients (40). We identified the irreversible ErbB1/2/4 inhibitor Poziotinib which was two orders of magnitude more cytotoxic in malignant BCBM94, with a single digit nanomolar IC50 compared to non-malignant HME1 cells **(Fig. 7A)**. Poziotinib half-maximal activity concentration **(AC_50_)** did not significantly differ between BCBM94 and BT474, indicating efficacy against both HER2+ BC models **(Fig. 7B)**. Furthermore, rhNRG1 failed to rescue HER2+ BC models from the cytotoxic effects of Poziotinib **(Fig. 7C)**. By contrast, the reversible RTKi Lapatinib and Tucatinib or the irreversible RTKi Neratinib only had moderate efficacy and rhNRG1 caused a shift in AC_50_, indicating that NRG1 successfully rescued BCBM94 from the cytotoxic ErbBi activity of these clinically used drugs **(Fig. 7C)**. Poziotinib emerged as a promising candidate for overcoming the anti apoptotic action of NRG1. Poziotinib (2.5-5nM) showed high cytotoxicity in cultured BCBM94 and BT474 cells **(Fig. 7D)** and rhNRG1 was unable to rescue these HER2+ BC cells as determined by WST **(Fig. 7E, F)** and cell impedance assays **(Sup. Fig. 3A, B)**. Poziotinib caused apoptosis with significant PARP cleavage **(Fig. 7G, H)** and, despite rhNRG1 present, abolished phosphorylation of ErbB3^Y1289^ **(Fig. 7I, J)** and Akt^S473^ **(Fig. 7K, L)**. We concluded that small molecule ErbB inhibitor Poziotinib blocked NRG1-ErbB3-PI3K-Akt signaling to cause apoptosis in HER2+ BC (**Fig.8**).

**Figure 7.**
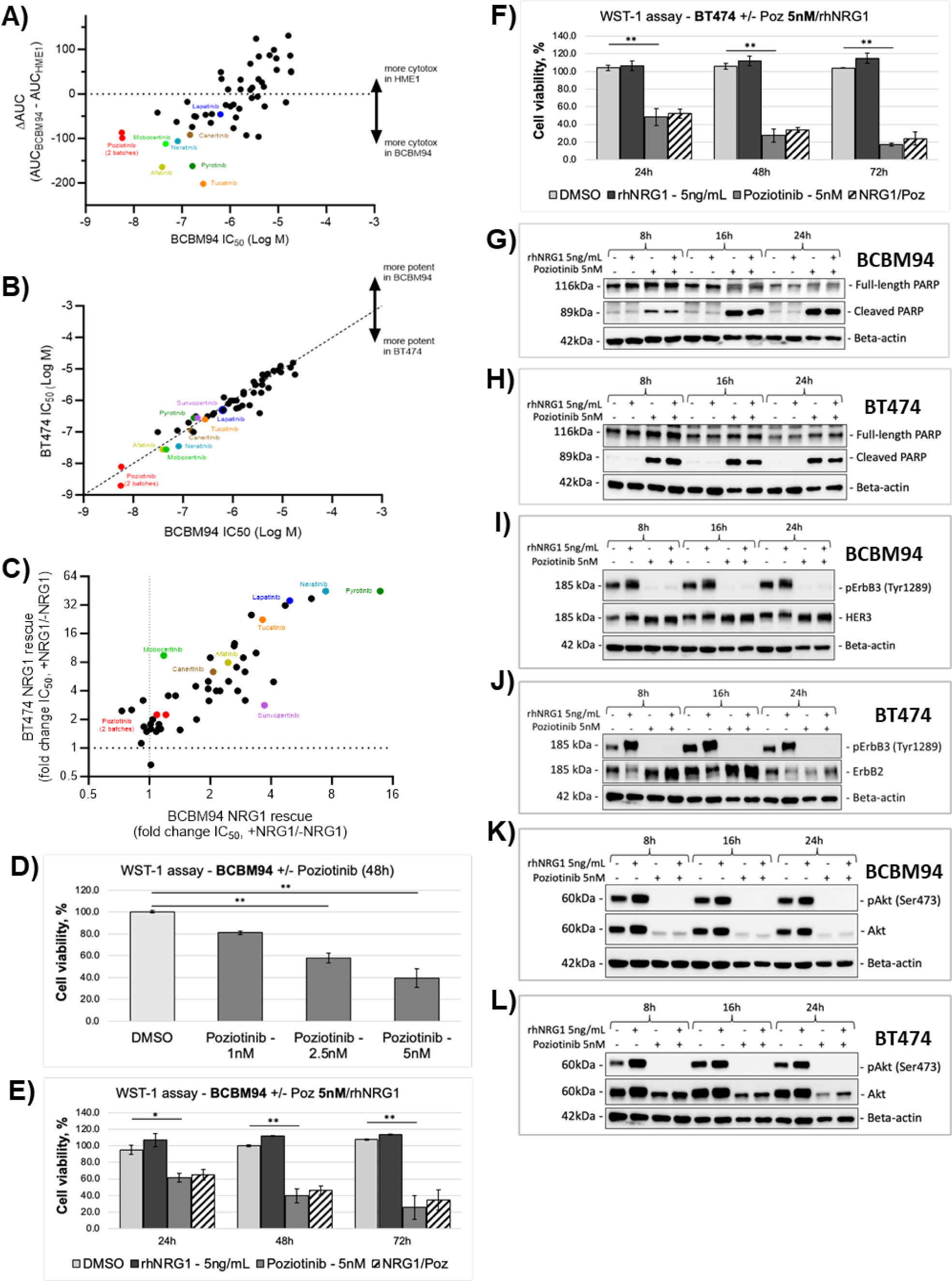
Poziotinib-induced cytotoxicity is not diminished by NRG1. **(A)** Fifty ErbB inhibitors were tested for differential sensitivity in BCBM94 versus HME1. 53 samples (3 compounds were represented by two separate batches) were tested at 22 concentrations ranging from 0.003pM to 40uM (BCBM94 n=4, HME1 n=1). The area under the curve (AUC, y-axis) of the non-linear fitting of the dose-response data was calculated as a metric to compare differential sensitivity to the compounds. The potency of each compound in the BCBM94 model is indicated on the x-axis. Compounds in the lower left corner of the plot are the most potent and BCBM94-selective. (**B)** Poziotinib is highly efficacious in reducing cell viability of both BCBM94 and BT474 HER2+ BC models. ErbB inhibitors were tested at concentrations ranging from 0.003 pM to 40uM and viability was measured using a CellTiterGlo assay after 72 h (BCBM94 n=4, BT474 n=1). (**C)** rhNRG1 reduces the cytotoxic activity of many ErbB inhibitors. The 50 ErbB inhibitors were tested +/- 5ng/mL rhNRG1 and viability was measured after 72h using CellTiterGlo. The differential sensitivity was assessed by comparing IC50 values (ratio) under both conditions (BCBM94 n=3, BT474 n=1). Data points that fall near the intersection of the dotted lines, including Poziotinib (Poz), represent compounds with equipotent activity +/- NRG1. Compounds to the upper right of the plot are those where NRG1 reduced the cytotoxic effect of the compound. (**D-F)** rhNRG1 failed to rescue tested HER2+ BC models from Poziotinib-mediated cytotoxicity. Cell viability of BCBM94 (n=3) **(D, E)** and BT474 (n=3) **(F)** under Poz +/-rhNRG1 treatment was assessed in the WST-1 assay. The endpoint absorbance readouts were used for quantification of the relative cell viability. (**G, H)** rhNRG1 failed to counteract Poziotinib-induced apoptosis. WB images show the levels of cleaved/ full-length PARP in BCBM94 (representative examples, n=3) (**G**) and BT474 (n=1) (**H)** cells under Poz +/-rhNRG1 treatment. (**I, J)** rhNRG1 was unable to rescue ErbB3 phosphorylation under Poziotinib. Representative WB images show the levels of phospho-/ total ErbB3 in BCBM94 (n=3) **(I)** and BT474 (n=1) **(J)** cells under Poz +/-rhNRG1 treatment. (**K, L)** rhNRG1 failed to rescue Akt phosphorylation under Poziotinib. Representative WB images show the levels of phospho-/ total ErbB3 in BCBM94 (n=3) **(K)** and BT474 (n=) **(L)** cells under Poz +/- rhNRG1 treatment. Bar charts present mean +/- SD, *p<0.05; **p<0.01.

**Figure 8.**
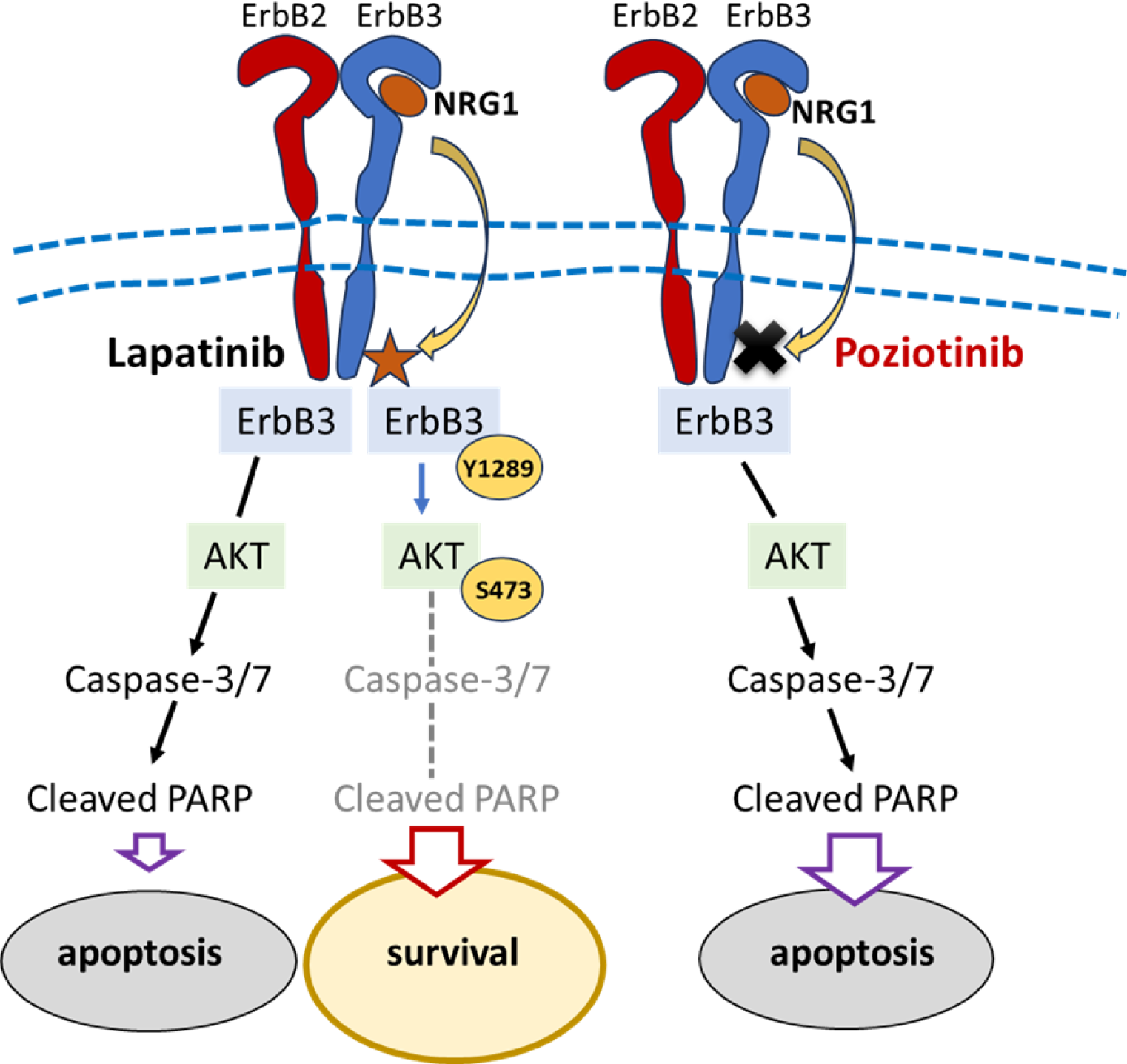
Schematic illustration showing the inability of NRG1 to rescue phosphorylation of ErbB3 and AKT resulting in PARP cleavage and Poziotinib-induced apoptosis in BCBM cells.

### Poziotinib eliminates HER2+ BC metastasis *in-vivo*

Computational predictions indicated high brain penetrability of Poziotinib which was confirmed in our pharmacokinetic studies in C57BL mice that demonstrated excellent brain permeability of Poziotinib when applied via two different routes. Our tolerability tests identified subcutaneous administration of Poziotinib to be far superior to oral application for subsequent drug treatments of BC brain metastases in C57BL mice (data not shown). Poziotinib administered at 5 mg/kg PO (per os) and SC (subcutaneous), and 2 mg/kg PO reached therapeutic concentrations in the brain, with concentrations of this RTKi still detectable at IC50 of 2.5 nM at 20h and 8h upon administration, respectively **(Suppl. Fig. 3C**).

To determine the *in-vivo* efficacy of Poziotinib towards HER2+ BC brain tumors, we orthotopically xenografted BCBM94 and BT474 cells into the right striatum of immunocompromised Rag2γc-/- mice. HER2+ BC brain tumors were confirmed by magnetic resonance imaging (MRI) prior to treating the animals with 100μl of either Lapatinib (80mg/kg, PO), Poziotinib (4mg/kg, SC), or solvent control (PO and SO) for two cycles of 5-days ON and 2 days OFF. MRI volumetry of pre- and post-treatment scans demonstrated a highly significant reduction in BCBM94 tumor volumes with Poziotinib, but not Lapatinib **(Fig. 9A).** The high efficacy of Poziotinib against HER2+ BC brain tumors was also confirmed in mice xenografted with BT474 cells **(Fig. 9B).** Ultra-performance liquid chromatography-tandem mass spectrometry (UPLC-MS/MS) analysis of plasma and brain samples collected on the last day of treatment (1h post-dosing) demonstrated that Poziotinib and Lapatinib reached similar concentrations in the brain at the doses administered. Notably, Poziotinib more effectively crossed the blood-brain barrier, with a two-fold higher brain/plasma concentration ratio than Lapatinib **(Fig. 9C)**. Post-treatment FFPE brain sections of mice xenografted with BCBM94 confirmed the observed MRI changes. BCBM94 tumors were exclusively identified in H&E stained tissues of the Lapatinib and solvent control groups. These tumors showed phosphorylated ErbB3^Y1289^ and Ki67+ nuclei and were largely negative in TUNEL tests detecting damaged DNA **(Fig. 9D)**. In sharp contrast, the tumor sites of all mice treated with Poziotinib were devoid of BCBM94 tumor cells, ErbB3 phosphorylated cells, and Ki67+ nuclei, but demonstrated positive TUNEL staining **(Fig. 9D)**. Similar results were obtained for mice orthotopically xenografted with BT474 cells and treated with Poziotinib or Lapatinib (data not shown).

**Figure 9.**
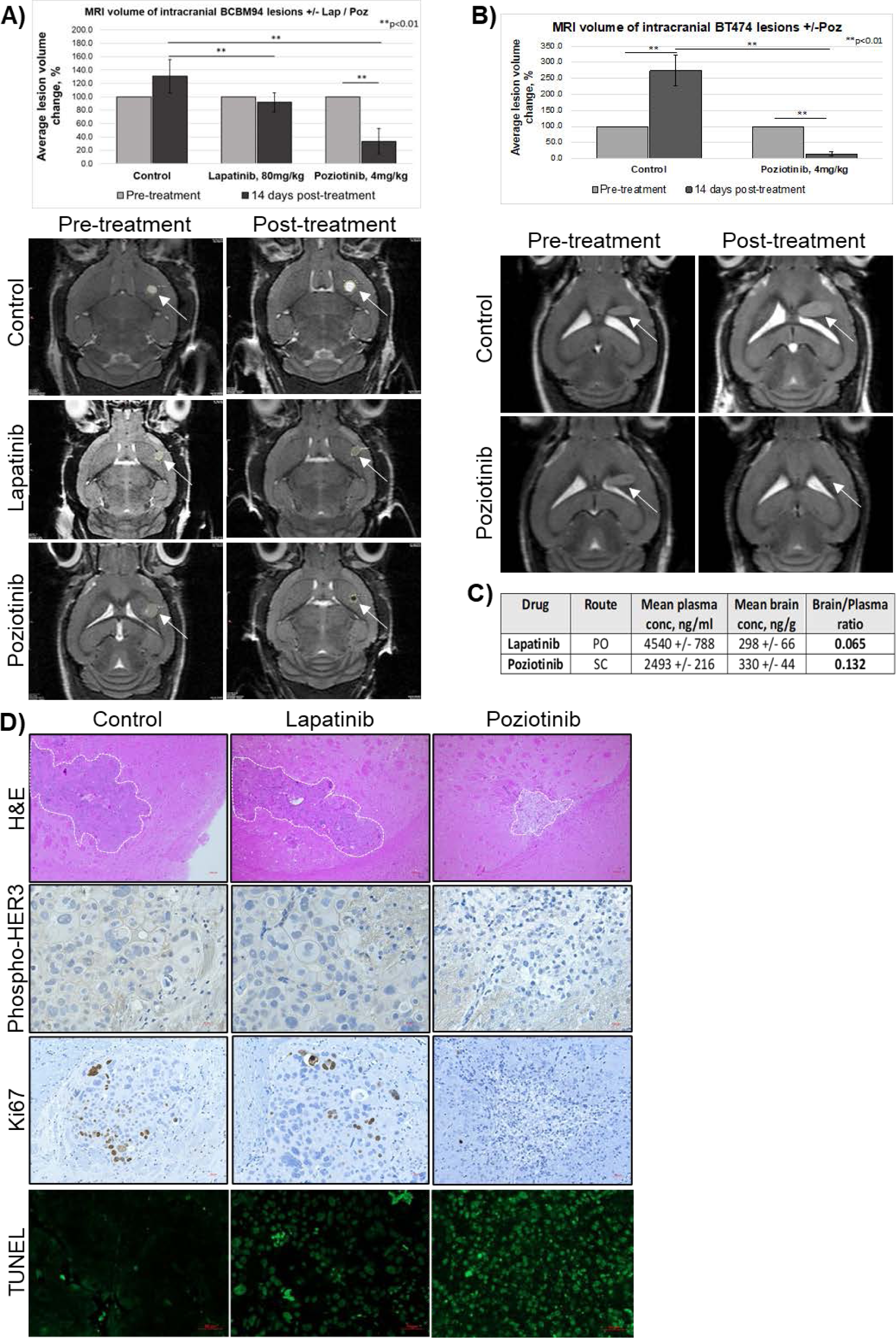
Poziotinib effectively reduced brain metastases in mice. BCBM94 and BT474 cells were orthotopically xenografted into SCID and RAG2yc-/- mice. Upon detection of sizable brain metastases with MRI, the animals were treated with either Lapatinib (80mg/kg), Poziotinib (4mg/kg), or solvent control for two 5-day cycles with 2 days off treatment in between. **(A)** Treatment with Poziotinib resulted in a significant reduction of BCBM94 and BT474 brain metastatic tumors. The ROI-based volumetry was performed on pre- and post-treatment sets of MRI scans (n=4/group). Bar charts present mean +/- SD, **p<0.01. Representative MR images are shown below the charts **(A, B)**. **(C)** Poziotinib is more brain penetrable than Lapatinib. Drug concentrations were measured by UPLC-MS/MS analysis of plasma and brain tissues taken 1 hour after the last drug administration (n=4/group). **(D)** Poziotinib abrogated ErbB3 phosphorylation, inhibited proliferation, and induced apoptosis within BCBM94 brain metastasis. H&E, IHC, and TUNEL analysis of post treatment mouse brain FFPE tissues is shown for solvent control, Lapatinib and Poziotinib treatment groups. The white dashed lines mark the margins of the metastatic lesion. ErbB3 phosphorylation and presence of Ki67+ nuclei were observed only in lesions of the solvent control and Lapatinib groups. TUNEL assay shows fragmented DNA in green color. Magnification: H&E 100x, Ki67 200x, TUNEL and pErbB3 400x.

## DISCUSSION

Despite successful therapies of peripheral BC disease and longer survival, patients with HER2+ BC have an increased risk of brain metastasis from HER2+ tumors (41, 42). Treatment with humanized monoclonal antibodies against HER2 and/or RTK small molecule inhibitors are not successful for HER2+ brain metastatic disease, in part because these compounds do not reach therapeutic concentrations in the brain (43-45). There is an urgent need for experimental *in-vivo* models to study HER2+ breast cancer brain metastasis because additional environmental factors in the brain metastatic niche can determine treatment responses. Among approximately 30 HER2+ BC cell lines that have been established so far, there are only a few xenogeneic (e.g. BT474) models that can cross the blood-brain barrier (BBB) and repeatedly establish hematogenous brain metastasis (46, 47). Although BT474 and our new BCBM94 model described here can be classified as a Luminal-B HER2+ molecular subtype, their receptor profile differs substantially. While BCBM94 is ErbB1-4-positive, ER-low, and PR negative, BT474 is ErbB1-4-positive, ER-high, and PR-positive. Another unique feature of BCBM94 is that, unlike BT474 which was isolated from primary breast cancer (46), BCBM94 is derived directly from a patient’s breast cancer brain metastasis. The molecular differences in both brain metastatic HER2+ BC models reflect a degree of molecular heterogeneity in BC brain metastasis. The ability of our BCBM94 model to cross the BBB and establish hematogenous brain metastasis combined with a unique receptor profile make the BCBM94 model a valuable new tool for studying multiple aspects of brain metastasis in HER2+ BC. Since BCBM94 require approximately 3.5 months to establish sizeable brain metastasis for hematogenic and orthotopic xenografting alike, our model reflects the long latency observed for human brain metastasis in HER2+ BC patients.

Upregulation of endogenous expression of NRG1 in several HER2+ BC cell lines (BT474, SKBR3) was previously shown to promote autocrine activation of the ErbB3-EGFR signaling axis and mediate resistance to Lapatinib (27). Our current data place exogenous NRG1 at the top of a powerful NRG1 ErbB3-PI3K-AKT-BAD signaling axis that mediates anti-apoptotic resistance against Lapatinib in the HER2+ BC metastatic brain niche. NRGs are abundantly expressed by resident cells of the normal brain (48, 49), as are ADAM metalloproteinases responsible for shedding of the extracellular EGF-like domains of membrane-bound NRG precursors (16, 17, 24, 25). We localized multiple NRG1+ cells in the TME of BCBM94 BC brain metastases as source for TME-derived NRG1 to induce therapeutic resistance to Lapatinib through paracrine and/or juxtacrine NRG1-ErbB3 signaling in brain metastases of HER2+ BC. Indeed, our in-vitro data revealed that exogenous NRG1 can counteract the cytotoxic and pro-apoptotic actions of Lapatinib in BCBM94 and BT474 cells.

The intrinsic apoptosis pathway (50) is primarily regulated by the Bcl-2 family of proteins consisting of the anti-apoptotic, pro-apoptotic BH3-only (apoptosis initiating), and pore-forming (executive) proteins (39) which initiate apoptosis through disruption of the outer mitochondrial membrane and the release of cytochrome C. In both HER2+ BCBM cell models, the apoptotic action of Lapatinib included a significantly reduced phosphorylation of ErbB1-4 and downstream targets AKT and BAD. Dephosphorylated BAD can engage in dimer formation with anti-apoptotic BCL-2 family members BCL-XL, BCL-2, and MCL-1, which prevents BAX/ BAK oligomeric pore formation in the outer mitochondrial membrane and the initiation of intrinsic caspase-mediated apoptosis (51). We confirmed BAX/ BAK oligomers by co-immunofluorescence and showed the induction of caspase 9 cleavage and activation of the effector caspases-3/7 in Lapatinib-treated BCBM94 and BT474 cells.

In agreement with clinical data, Lapatinib was ineffective in reducing the growth of HER2+ BC tumors in mouse brain, despite reaching concentrations that reduced viability in-vitro. Our results suggest that the presence of brain-derived NRG1, which could compete with Lapatinib for binding to ErbB3/4 expressed on both BCBM94 and BT474 BC cells, may account for endocrine therapeutic resistance observed in our mouse studies and in the clinic. To be considered a promising candidate for the treatment of HER2+ BC brain metastases, any of the 50 small molecule ErbB inhibitor compounds tested in our high-throughput screens had to meet stringent selection criteria. This included good brain permeability, high efficacy at low nanomolar concentrations, selective toxicity to HER2+ BC cells but not to non tumor breast epithelial cell line HME-1, and the ability to inhibit the growth of our HER2+ BC cell models in the presence of exogenous NRG1. Of several potential candidates, the irreversible ErbB inhibitor Poziotinib had the lowest IC_50_ (low nanomolar) and showed similar efficacy in both BCBM94 and BT474 models but had minimal toxicity to HME-1. A powerful inducer of apoptosis, irreversible RTKi Poziotinib blocked rhNRG1-mediated rescue of the ErbB3-AKT-Bad signaling cascade in both tested HER2+ brain metastatic BC models. BCBM94 and BT474 xenografts demonstrated a dramatic reduction in tumor volume after only two weeks of treatment with Poziotinib, whereas xenografts continued to grow under Lapatinib. In agreement with a significant tumor volume reduction on MRI scans, histological examination demonstrated the absence of viable tumor cells, the loss of ErbB3 phosphorylation, and marked DNA fragmentation at brain injection sites upon Poziotinib treatment. By contrast, BCBM94 lesions treated with Lapatinib remained proliferative and had preserved ErbB3 phosphorylation. Poziotinib, alone or in combination with Fulvestrant, was recently shown to attenuate tumor growth, multiorgan metastasis, and mTOR activation in recurrent metastasizing BC cells harboring an HER2^L755S^ mutation that conferred resistance to the irreversible ErbB inhibitor Neratinib. Recent clinical trial phase I and II studies used orally administered Poziotinib in patients with metastasising breast cancer (NOV120101-203 trial) (52) and non-small cell lung cancer with HER2 exon 20 insertions (ZENITH20-2 Trial) (53, 54). Although Poziotinib showed meaningful clinical activity in these heavily pretreated HER2+ metastatic BC and non-small cell lung cancer patients, including a small group of patients with brain metastases, the toxic side effects, mainly rash, diarrhea, and stomatitis, frequently led to dose reductions. In our animal studies, we noticed that the same concentration of Poziotinib proven effective at treating BCBM94 and BT474 xenografts showed significantly higher toxicity upon oral than subcutaneous administration in mice. Our pharmacokinetic studies confirmed that either administration route yielded similar brain concentrations, suggesting a possible remedy to the clinical toxicities observed with Poziotinib in patients. Recently, the irreversible ErbB1/2 kinase inhibitor Pyrotinib showed beneficial effects in patients with advanced metastatic HER2+ BC who had progressed under Trastuzumab (55). For patients with radiotherapy-naïve and radiotherapy resistant HER2+ brain metastases, Pyrotinib in combination with capecitabine was reported to have a response rate of 74% and 42%, respectively (55). However, our small molecule compound screens revealed that the AC_50_ for Pyrotinib required 10 times higher concentrations than Poziotinib. Importantly and in contrast to Poziotinib, the efficacy of Pyrotinib was reduced 10 fold in the presence of NRG1 in both of our HER2+ BCBM cell models (Suppl. Table 1).

In summary, we have established and characterized a novel patient-derived brain metastatic HER2+ BC cell line capable of hematogenic colonization of mouse brain as pre-clinical model to study HER2+ BC brain metastasis. We unveiled the antiapoptotic mechanism of exogenous NRG1 as a driver of resistance to the reversible RTKi Lapatinib in HER2+ BC brain metastases. Poziotinib prevented the anti-apoptotic actions of NRG1 in-vitro and revealed high efficacy in abrogating HER2+ BC brain metastatic lesions in mice. When administered by SC route, this irreversible RTKi is a promising drug for the management of brain metastases of HER2+ BC.

## LIMITATIONS

We confirmed the ability of BCBM94 to establish hematogenous brain metastasis after intracardial xenografting. However, we performed orthotopic intracranial, and not intracardial, xenografting of HER2+ BC cell in mice when testing the effects of Lapatinib and Poziotinib in-vivo in order to measure the size of lesions at identical brain locations during treatments. Future studies will use intracardial xenografting to determine if hematogenous HER2+ BC brain metastases undergo similar regression with Poziotinib.

## MATERIALS AND METHODS

### Cell Culture

Human HER2+ BT474 (HTB-30) and SKBR3 (HTB-20) BC cell lines were acquired from ATCC. The brain-seeking subline of the triple-negative MDA-MB-231 cells, MDA-MB-231/BR, was a generous gift of Dr. Patricia Steeg (NCI, Bethesda, Maryland). A patient-derived HER2+ BCBM94 BC cell line was established in the laboratory from BC cells isolated from a surgically resected HER2+ BC brain metastasis. The first mouse brain passage of the BCBM94 model was used in the study. All cell lines were routinely cultured in DMEM/F-12 Ham’s medium supplemented with 10% FBS. Except for data presented in **Fig. 6A-C** in which 10 % FBS was supplemented during the assay, all treatments were done in DMEM/F-12 Ham’s 1% FBS. hTERT-HME1 (CRL-4010) were acquired from ATCC and cultured in Mammary Epithelial Cell Growth Medium Bullet Kit from Lonza (Bend, OR, Catalog #: CC-3150).

### siRNA gene silencing

ErbB3 silencing was achieved with 20nM of ErbB3 siRNA purchased from Ambion (Cat. AM16708, ID146247). The non-coding siRNA was acquired from Dharmacon (Horizon, St. Louis, MO, Cat. D-001810-10-20). siLentFect lipid reagent (BioRad, Mississauga, ON, Cat. 1703360) was utilized for transfection.

### Drugs and growth factors

Reversible EGFR/ErbB2 RTK inhibitor Lapatinib (Cat. S1028), irreversible pan-ErbB RTK inhibitor Poziotinib (Cat. S7358), and PI3K/Akt/mTOR inhibitor PI-103 (Cat. S1038) were purchased from Selleckchem (Houston, TX). Recombinant human NRG1 was acquired from BioLegend (San Diego, CA, Cat. 551904). High-throughput drug screening assays were used with compounds sourced as indicated in **Suppl. Table 1**.

### Blood brain barrier penetrability predictions

CNS penetrance was predicted by taking a consensus of the predictions derived from the following methods: 1. The Blood–Brain Barrier (BBB) Score (56), using a cutoff >=3.5 to designate a compound as CNS penetrant; 2. B3clf Predictors for Blood-Brain Barrier Permeability with resampling strategies based on B3DB database (57); 3. Using the BBB permeability classification model (BBB-Filter) from ADMET Predictor software (58).

### Cell based assays

Cell proliferation reagent WST1 was acquired from Sigma Millipore (Cat. 5015944001) and used as per the manufacturer’s protocol. BCBM94 and BT474 were seeded at 4000 cells/well in 100uL volume in a 96-well plate. After 24h the culture medium was replaced with 100uL/well media containing treatment compounds. Caspase-Glo 3/7 Assay System (Promega, Cat. G8091) was used to measure the activity of caspase-3 and caspase-7. Cell seeding and incubation conditions were identical to that described for the WST1 assay. For Colony formation assays (59), BCBM94 were seeded at 20,000 cells/well in 6-well plates. The culture media was replaced every 3 days. After 14 days the cells were fixed and imaged with a D2 inverted microscope (Zeiss, Jena, Germany). For the high-throughput drug screening assay, compounds were added to 1536-well white-walled plates (Greiner, Monroe, NC, Cat. 789173) using acoustic dispensing. Cells were then seeded at 500 cells/well in 5uL volume. After 72h of incubation (48h for HME1 assay), 2.5uL of CellTiterGlo reagent (Promega, Madison, WI, Cat. G7570) was added to each well and plates were incubated at room temperature for 10 mins prior to luminescence readout. For Mitotracker assays, BCBM94 cells were seeded at 80,000 cells/well on glass coverslips in 6-well plates. At the end of treatments, MitoTracker Red CMXRos (Cell Signaling Technology, Danvers, MA, Cat. 9082) was added to a final concentration of 100 nM/ well and plates were incubated for 30 min at 37°C in the dark. Cells were fixed with ice-cold methanol and the mean fluorescence intensity was quantified per 100,000 µm² and 200 nuclei per treatment group using Zen software (Zeiss).

### Quantitative real-time polymerase chain reaction (qRT-RCR)

Total RNA was isolated with TRIZol and cDNA was synthesized with qScript cDNA SuperMix (QuantaBio, Beverly, MA, Cat. 95048-025). Primer sequences are listed below and the SYBR Green PCR Master Mix (Applied Biosystems, Cat. 4300155) was used. mRNA expression was analyzed by QuantStudio 3 qRT-PCR System (Applied Biosystems, Fisher Scientific, Winnipeg, MB), and the Comparative CT (ΔΔCt) Method (60). Primer sequences were: F_huNRG1, 5’ ATTGAAAGAGATGAAAAGCCAGG-3’; R_huNRG1, 3’-GCCAGTGATGCTTTGTTAATGC-5’; F_huNRG2, 5’-CTAAGCAAAAAGCCGAGGAGC-3’; R_huNRG2, 3’ CTTCCGCTGTTTTTTGGTCTTG-5’; F_huNRG3, 5’ AACACTTATCATTGGAGCCTTC-3’; R_huNRG3, 3’-GGTGTTTCATTTTCTGCCTTTG-5’; F_huNRG4, 5’ CTCTGGGTATTGTGTTGGCTG-3’; R_huNRG4, 3’ TGTCCTCCTGCACCAAAAACC-5’.

### Gel electrophoresis of proteins and Western blotting

The whole-cell lysates prepared with Laemmli cell lysis buffer were separated on TGX FastCast acrylamide gels (BioRad, Mississauga, ON). Nitrocellulose membranes and the Trans-Blot Turbo system (Bio-Rad) were used for protein transfer. The non-specific antibody binding sites were blocked with 5% non-fat milk in TBST, pH 7.6 for 1h at RT. Antibodies and incubation conditions are listed in **Table 1**. Proteins were visualized using Clarity (Max) Western ECL Substrates (Bio-Rad, Cat. 1705060, Cat. 1705062) and ChemiDoc MP Imaging System (Bio-Rad). Band volume quantification (densitometry) was done in the Image Lab (Bio-Rad).

**Table 1:**
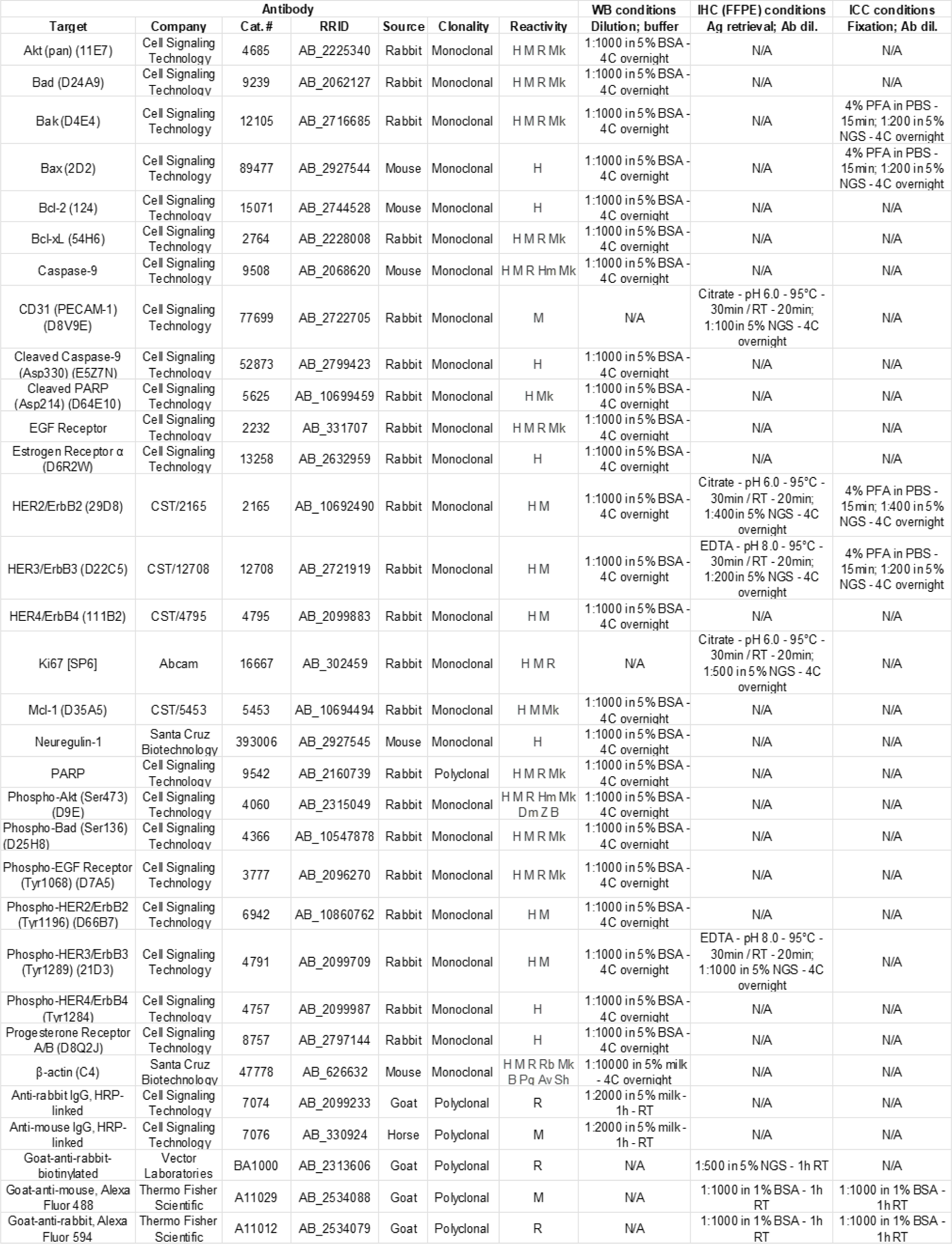
antibodies used in this study.

### In situ hybridization

Formalin-fixed, paraffin-embedded (FFPE) mouse brain tissue slides were processed according to the manufacturer’s instructions (61). Hybridization probes against human ErbB2 (Cat. 418741) and mouse Nrg1 (Cat. 441811) were acquired from ACDBio (Newark, CA). Hybridization, amplification, and signal detection were performed according to the manufacturer’s protocol (62). Signal amplification and detection were performed with RNAscope 2.5 HD Duplex Reagent Kit (ACDBio, Cat. 322430).

### Immunodetection in brain tissues and cells

Mouse FFPE brain tissue sections were deparaffinized and re-hydrated. The list of antibodies, antigen retrieval, and staining conditions are listed in **Table 1**. Non-specific antibody binding sites were blocked with 10% normal goat serum in TBST pH 7.6 for 1h. The chromogenic signal was amplified using HRP-Streptavidin (Jackson ImmunoResearch, West Grove, PA, Cat. 016-030-082) and detected with DAB Substrate Kit (Fisher Scientific, Winnipeg, MB, Cat. PI34002). BCBM94 and BT474 were seeded at 80,000 cells/well on glass coverslips in 6-well plates. Cells were fixed with 4% methanol-free formaldehyde in PBS for 15 min at room temperature (RT) and permeabilized in 0.1% Triton X-100 in PBS for 15 min at RT. Non-specific antibody binding sites were blocked with 5% NGS in PBS for 1h. For antibodies and staining conditions see **Table 1**.

### TUNEL assay

In Situ Cell Death Detection Kit, POD (Roche, Mississauga, ON, Cat. 11684817910) was used as per manufacturer’s protocol to detect apoptosis in FFPE mouse brain tissue sections.

### Transmission electron microscopy (TEM)

BCBM94 cells were trypsinized, pelleted, and fixed in 3% Glutaraldehyde in 0.1M Sorensen’s buffer and post-fixed in 1% Osmium tetroxide (OsO4) in 0.1M Sorensen’s buffer. The pellets were embedded in EMBed 812 resin (Electron Microscopy Sciences, Hatfield, PA, Cat. 14900). Thin sections were stained with Uranyless (EMS, Cat. 2240920) and Lead citrate (EMS, Cat. 22410) and imaged using Philips TM10 transmission electron microscope. At least 10 cells per sample group were imaged and qualitatively analyzed for signs of mitochondrial damage.

### *In-vivo* experiments

Ultrasound-guided intracardial injection of BCBM94 cells into RAG2γc-/- mice (50) resulted in brain metastatic tumors after around 3.5 months. The animal research was approved by the Bannatyne Campus Animal Care Committee (protocol #21-017). We opted for orthotopic xenografting into the right striatum using stereotactic surgery to achieve the same lesion size at the identical brain location for BCBM94 and BT474 tumor growth in mouse brain (63). Intracranial tumor growth was monitored using MRI. Upon detection of sizeable brain metastases, the animals were treated with either Lapatinib (PO, 80mg/kg) or Poziotinib (SC, 4mg/kg) for two cycles of 5 days ON and 2 days OFF. Solvent controls were 10% solutol in water (for Poziotinib) and Phosal/PEG300 (for Lapatinib). MR images were acquired using a 7T cryogen-free superconducting magnet (MR Solutions, Boston, MA, USA) with a 17cm bore and equipped with a dedicated quadrature mouse head coil. Animals were anesthetized with isoflurane and transferred to a warmed Minerve© bed for head fixation and further anesthetic delivery. Respiration rates were monitored throughout the procedure. Following a preliminary scout scan, acquisition planes were optimized, and whole-brain coronal T2-weighted scans were acquired using a fast spin echo sequence with the following parameters: TR 5000ms, TE 45ms, echo trains 7, FOV 30x30mm^2^, matrix size 250x256, total slices 18, slice thickness 0.3mm, and two averages for a total scan time 350s. For MRI lesion volumetry, region of interest (ROI)-based volumetry was performed on pre- and post-treatment sets of MRI scans (n=4/ group). The total MRI lesion volume was quantified as a sum of individual volumes (V = πr^2^h) for each MRI section containing the lesion. Measurement of drug penetration into mouse brain were performed with C57BL mice. Poziotinib was administered by either oral gavage (PO) or subcutaneous injection (SC). Plasma and brain tissues were collected immediately after the last treatment and snap-frozen in liquid nitrogen for ultra-performance liquid chromatography-tandem mass spectrometry (UPLC-MS/MS) analysis. The calibration standards and quality control samples were prepared in the blank mouse plasma and brain homogenate. Aliquots of 10 µL samples were mixed with 200 µL internal standard in acetonitrile to precipitate proteins in a 96-well plate. 1.0 µL supernatant was injected for the UPLC-MS/MS analysis. MassLynx and TargetLynx were used for data collection and processing (Waters Corp., Milford, MA). For brain histology and IHC, mice were euthanized using isofluorane and cervical dislocation upon two treatment cycles with 5 days ON and 2 days OFF. Brains were removed immediately and hemispheres were fixed in 10% neutral buffered formalin for 24h. FFPE brain sections of 5µm were subjected to H&E staining and IHC.

### Statistical analysis

A paired two-tailed t-test was used to determine if there was a significant difference between the means of the two groups. One-way ANOVA was used to identify any statistically significant difference between the means of three or more groups. The Tukey test was performed to check if there is a statistically significant difference between the pairs of samples that belong to a larger group where the statistical difference proved to be significant in the ANOVA test.

## ACKNOWLEDGEMENTS

The brain-seeking subline of the MDA-MB-231 cell line, MDA-MB-231/BR, was kindly provided by Dr. Patricia Steeg (NCI, Bethesda, Maryland). We thank the nurses Deb Swan, Coleen Unger and Susan Pearce at the Neurosurgery Clinic, Health Sciences Center, Winnipeg, for their tremendous support with patient consent. The authors acknowledge the support from the staff at the Electron Microscopy Core Platform and the Histology Services, Department of Human Anatomy and Cell Sciences, University of Manitoba. We acknowledge the staff at the Central Animal Care Services and the Small Animal Imaging Core Facility, University of Manitoba, for their help throughout the study. We acknowledge funding support from the Natural Sciences and Engineering Research Council of Canada (TK), CancerCare Manitoba (TK, SHK), the Dr. Paul H.T. Thorlakson Foundation Fund (SHK), the Cancer Research Society (SHK), Mitacs (DI), the Max Rady College of Medicine (TT), the University of Manitoba Graduate Fellowship (JS), and the NCATS/NIH Intramural Research Program (YHL, XX, AW, RC, AK, JJM, MJH).

## Author contributions

Conceptualization of study (TK, SHK); data curation and formal analysis (DI, YHL, AG, TT, AK); animal surgeries (TK, JS); animal imaging (JS); patient neurosurgery (JB), pathology diagnosis (MDB); drug formulation (RC); drug screening methodology (YHL, MJH), pharmacokinetic data (AW, XX); resources and supervision (JJM, MJH, SHK, TK).

## SUPPLEMENTARY FIGURES

**Suppl. Figure 1:**
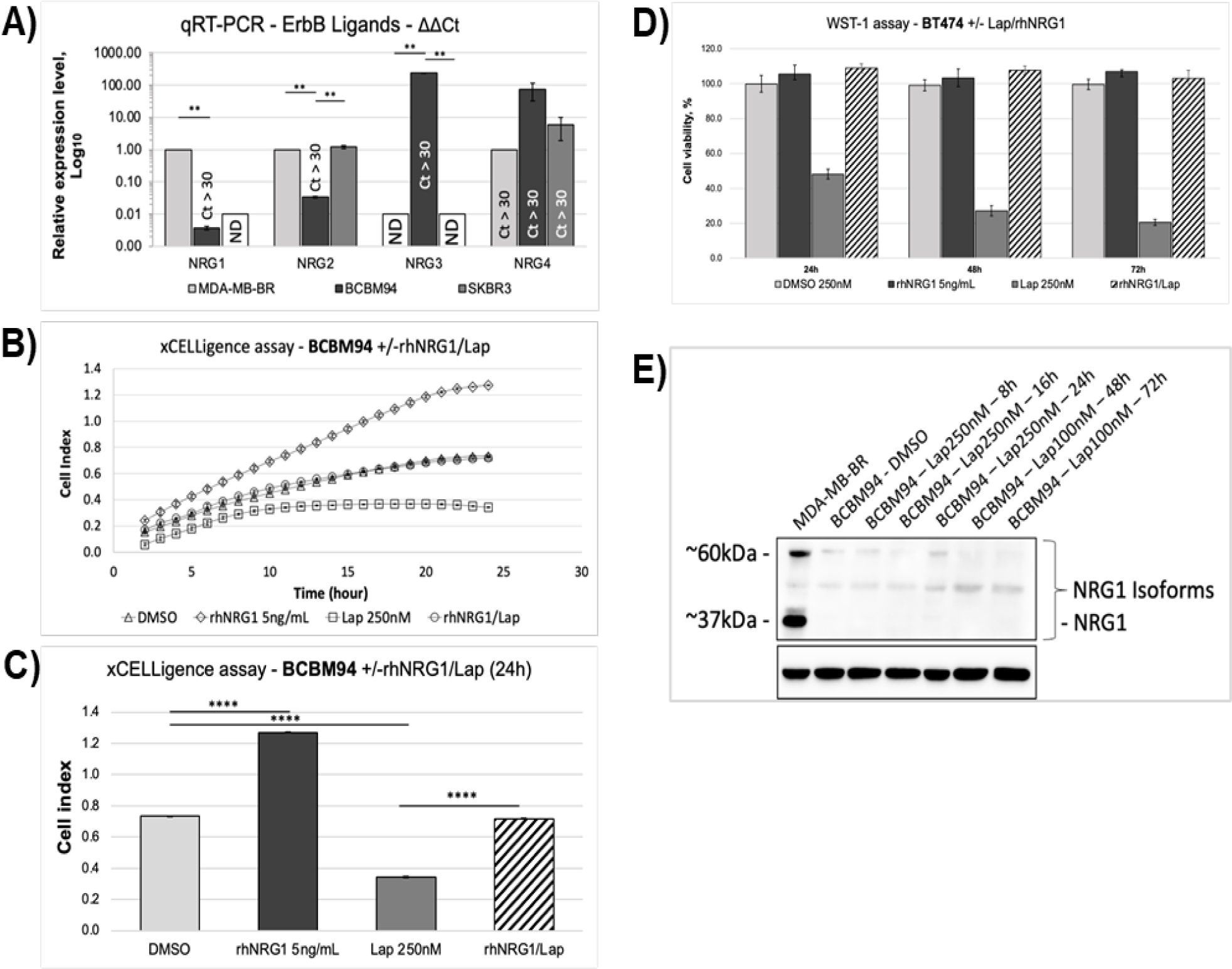
NRG1 counteracts Lapatinib-induced loss of viability A) BCBM94 cells have very low expression of NRG1-4 mRNA in-vitro. qRT-PCR compared mRNA levels of NRG1-4 in the triple-negative MDA-MB-BR and HER2+ SKBR3 and BCBM94 cells. A cycle count higher than 30 (Ct > 30) is considered as low expression independent of relative values. Values for MDA-MB-BR are set to 1. To calculate the relative expression of NRG3 in BCBM94, Ct values for MDA-MB-BR and SKBR3 were set to 40 (maximum cycle number). **ND**: not detected. **B,C) rhNRG1 rescued BCBM94 cells from Lapatinib-induced cytotoxicity.** Real-time cell analysis based on impedance measurements was performed with the xCELLigence assay. The graph depicts measurements of BCBM94 cells treated with Lap +/-rhNRG1 over a period of 25 hours **(B)**. The terminal 24h readouts of the xCELLigence assay were plotted on the bar graph **(C)**. **D) rhNRG1 rescued BT474 cells from Lapatinib-induced cytotoxicity.** Cell viability of BT474 cells under Lap +/- rhNRG1 treatment was assessed in the WST-1 assay. The endpoint absorbance readouts were used for quantification of the relative cell viability. **E) Lapatinib did not upregulate endogenous expression of NRG1 in BCBM94 cells.** WB images present the level of NRG1 protein in BCBM94 cells under two different concentrations of Lapatinib. MD-MB-BR cells serve as positive control for NRG1.

**Suppl. Figure 2:**
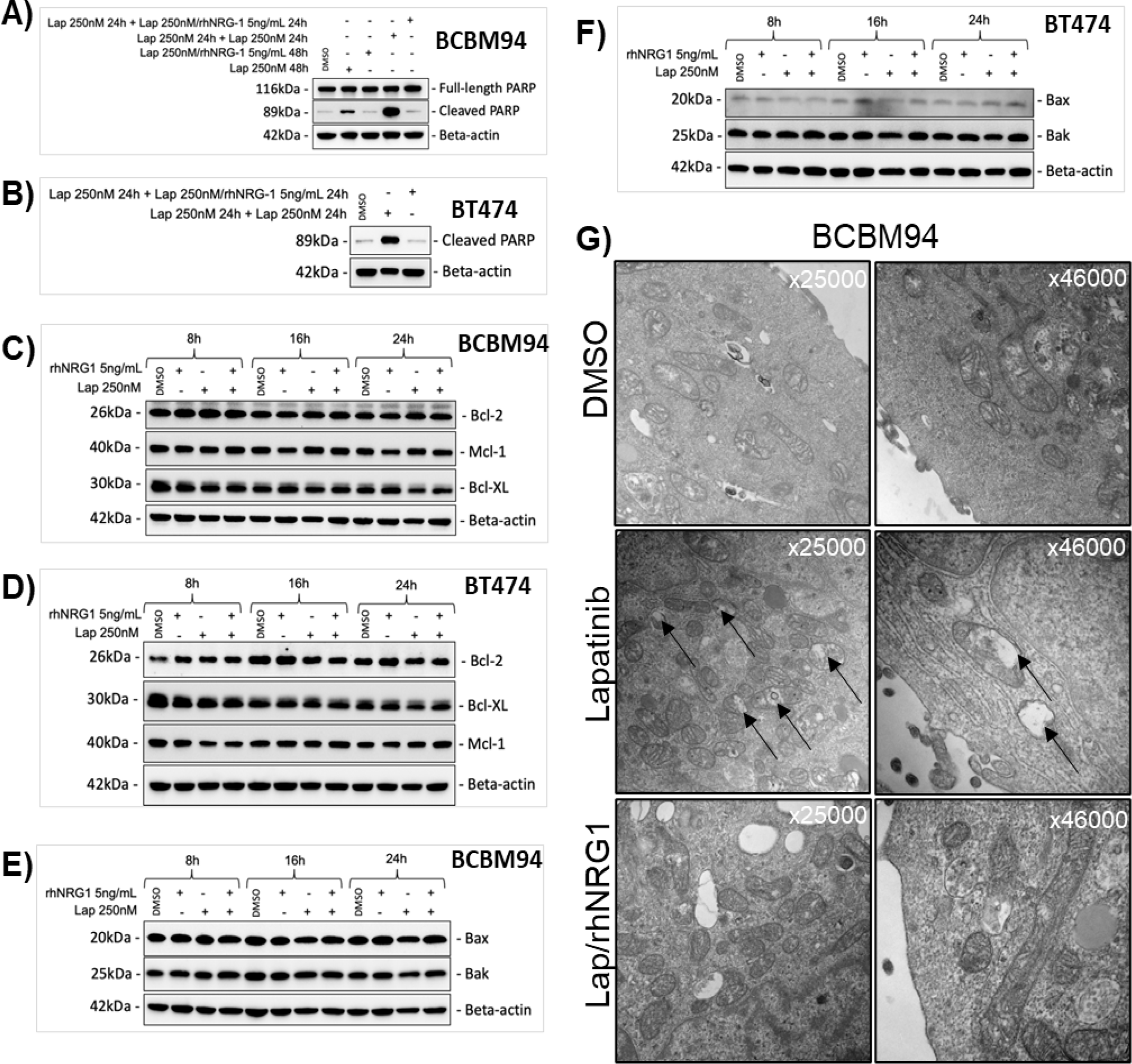
NRG1 counteracts Lapatinib-induced apoptosis A, B) rhNRG1 rescued BCBM94 and BT474 cells from Lapatinib-induced apoptosis even after pre-incubation with Lapatinib. BCBM94 and BT474 cells were treated with Lapatinib (Lap) for the first 24h and either Lapatinib or Lap/ rhNRG1 for another 24h. WB images show the levels of cleaved and full-length PARP proteins in BCBM94 **(A)** and BT474 **(B)** cells under the above-mentioned conditions. **C-F) Total levels of the antiapoptotic and proapoptotic (effectors) proteins of the Bcl-2 family were unaffected by Lap +/-rhNRG1 treatment.** WB images present the levels of Bcl-2, Bcl- XL, Mcl-1, Bax and Bak protein in BCBM94 **(C,E)** and **BT474 (D,F)** cell lines under Lap +/-rhNRG1 treatment for the indicated times. **G) Transmission electron microscopy (TEM) images.** NRG1 maintained mitochondrial cristae structure under Lapatinib. TEM images of BCBM94 cells incubated with Lap +/-rhNRG1. Black arrows indicate damaged mitochondria under Lapatinib treatment. Magnifications are indicated on the image.

**Suppl. Figure 3:**
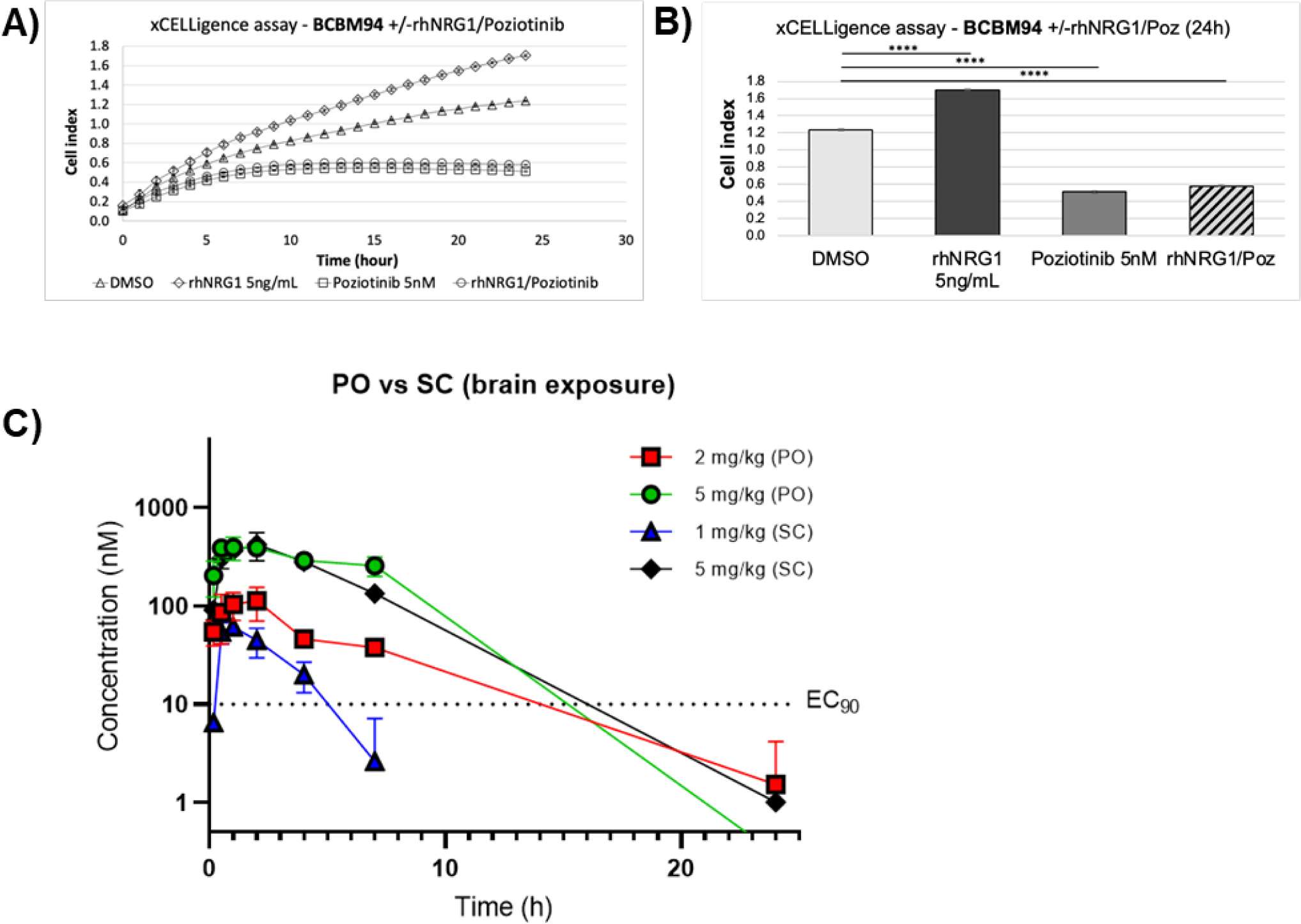
NRG1 fails to rescue poziotinib-induced viability loss. A, B) rhNRG1 failed to rescue BCBM94 cells from Poziotinib (Poz)-induced cytotoxicity. Real-time cell analysis using the xCELLigence assay show cell impedance measurements in BCBM94 cells treated with Poz +/-rhNRG1 over a period of 25 hours **(A)**. The terminal 24h readouts of the xCELLigence assay were plotted on the bar graph **(B)**. Bar charts present mean +/- SD, **p<0.01; ***p<0.001; ****p<0.0001. **C) Poziotinib brain concentrations.** Poziotinib was administered in C57BL/6 mice by oral gavage (PO) at 2 and 5 mg/kg and by subcutaneous injections (SC) at 1 and 5 mg/kg. Concentrations in brain tissues were determined by UPLC-MS/MS bioanalytical method at different time points after drug administration. The dotted line represents the IC_90_ concentration, as determined by in-vitro viability experiments.

## REFERENCES

1. The-American-Cancer-Society. Treatment of Breast Cancer By Stage. 2022.

2. Leone J, Lin N. Systemic Therapy of Central Nervous System Metastases of Breast Cancer. Current oncology reports. 2019;21(6).

3. Schlam I, Swain SM. HER2-positive breast cancer and tyrosine kinase inhibitors: the time is now. NPJ Breast Cancer. 2021;7(1):56.

4. National-Cancer-Institute. Cancer Treatment. 2020.

5. Zhong L, Y L, Xiong L, Wang W, Wu M, Yuan T, et al. Small molecules in targeted cancer therapy: advances, challenges, and future perspectives. Signal transduction and targeted therapy. 2021;6(1).

6. Lin N, Carey L, Liu M, Younger J, Come S, Ewend M, et al. Phase II trial of lapatinib for brain metastases in patients with human epidermal growth factor receptor 2-positive breast cancer. Journal of clinical oncology : official journal of the American Society of Clinical Oncology. 2008;26(12).

7. Lin N, Diéras V, Paul D, Lossignol D, Christodoulou C, Stemmler H, et al. Multicenter phase II study of lapatinib in patients with brain metastases from HER2-positive breast cancer. Clinical cancer research : an official journal of the American Association for Cancer Research. 2009;15(4).

8. Freedman R, Gelman R, Anders C, Melisko M, Parsons H, Cropp A, et al. TBCRC 022: A Phase II Trial of Neratinib and Capecitabine for Patients With Human Epidermal Growth Factor Receptor 2 Positive Breast Cancer and Brain Metastases. Journal of clinical oncology : official journal of the American Society of Clinical Oncology. 2019;37(13).

9. Bachelot T, Romieu G, Campone M, Diéras V, Cropet C, Dalenc F, et al. Lapatinib plus capecitabine in patients with previously untreated brain metastases from HER2-positive metastatic breast cancer (LANDSCAPE): a single-group phase 2 study. The Lancet Oncology. 2013;14(1).

10. Ullrich A, Schlessinger J. Signal transduction by receptors with tyrosine kinase activity. Cell. 1990;61(2).

11. Citri A, Skaria K, Yarden Y. The deaf and the dumb: the biology of ErbB-2 and ErbB-3. Experimental cell research. 2003;284(1).

12. Roskoski R. The ErbB/HER family of protein-tyrosine kinases and cancer. Pharmacological research. 2014;79.

13. Weitsman G, Barber PR, Nguyen LK, Lawler K, Patel G, Woodman N, et al. HER2-HER3 dimer quantification by FLIM-FRET predicts breast cancer metastatic relapse independently of HER2 IHC status. Oncotarget. 2016;7(32):51012–26.

14. Berghoff AS, Bartsch R, Preusser M, Ricken G, Steger GG, Bago-Horvath Z, et al. Co overexpression of HER2/HER3 is a predictor of impaired survival in breast cancer patients. Breast. 2014;23(5):637–43.

15. Jia R, Zhao H, Wang S. Neuregulin Signaling in the Tumor Microenvironment. Advances in experimental medicine and biology. 2021;1270.

16. Luo X, Prior M, He W, Hu X, Tang X, Shen W, et al. Cleavage of neuregulin-1 by BACE1 or ADAM10 protein produces differential effects on myelination. The Journal of biological chemistry. 2011;286(27).

17. Fleck D, van Bebber F, Colombo A, Galante C, Schwenk B, Rabe L, et al. Dual cleavage of neuregulin 1 type III by BACE1 and ADAM17 liberates its EGF-like domain and allows paracrine signaling. The Journal of neuroscience : the official journal of the Society for Neuroscience. 2013;33(18).

18. Wieduwilt M, Moasser M. The epidermal growth factor receptor family: biology driving targeted therapeutics. Cellular and molecular life sciences : CMLS. 2008;65(10).

19. Li Q, Ahmed S, Loeb J. Development of an autocrine neuregulin signaling loop with malignant transformation of human breast epithelial cells. Cancer research. 2004;64(19).

20. Cabrera R, Mao S, Surve C, Condeelis J, Segall J. A novel neuregulin - jagged1 paracrine loop in breast cancer transendothelial migration. Breast cancer research : BCR. 2018;20(1).

21. Jeong H, Kim J, Lee Y, Seo J, Hong S, Kim A. Neuregulin-1 induces cancer stem cell characteristics in breast cancer cell lines. Oncology reports. 2014;32(3).

22. The Human Protein Atlas. 2022(21.1).

23. Law AJ, Shannon Weickert C, Hyde TM, Kleinman JE, Harrison PJ. Neuregulin-1 (NRG-1) mRNA and protein in the adult human brain. Neuroscience. 2004;127(1):125–36.

24. Guo Z, Su Y, Wang Y, Wang W, Guo D. The expression pattern of Adam10 in the central nervous system of adult mice: Detection by in situ hybridization combined with immunohistochemistry staining. Molecular medicine reports. 2016;14(3).

25. Dominguez-Garcia S, Castro C, Geribaldi-Doldán N. ADAM17/TACE: a key molecule in brain injury regeneration. Neural regeneration research. 2019;14(8).

26. Valiente M, Van Swearingen AED, Anders CK, Bairoch A, Boire A, Bos PD, et al. Brain Metastasis Cell Lines Panel: A Public Resource of Organotropic Cell Lines. Cancer Res. 2020;80(20):4314–23.

27. Xia W, Petricoin E, Zhao S, Liu L, Osada T, Cheng Q, et al. An heregulin-EGFR-HER3 autocrine signaling axis can mediate acquired lapatinib resistance in HER2+ breast cancer models. Breast cancer research : BCR. 2013;15(5).

28. D’Amato V, Raimondo L, Formisano L, Giuliano M, De Placido S, Rosa R, et al. Mechanisms of lapatinib resistance in HER2-driven breast cancer. Cancer treatment reviews. 2015;41(10).

29. Garrett J, Olivares M, Rinehart C, Granja-Ingram N, Sánchez V, Chakrabarty A, et al. Transcriptional and posttranslational up-regulation of HER3 (ErbB3) compensates for inhibition of the HER2 tyrosine kinase. Proceedings of the National Academy of Sciences of the United States of America. 2011;108(12).

30. Hegde P, Rusnak D, Bertiaux M, Alligood K, Strum J, Gagnon R, et al. Delineation of molecular mechanisms of sensitivity to lapatinib in breast cancer cell lines using global gene expression profiles. Molecular cancer therapeutics. 2007;6(5).

31. SW B, J Z, MH T, D Y. PI3K-independent mTOR activation promotes lapatinib resistance and IAP expression that can be effectively reversed by mTOR and Hsp90 inhibition. Cancer biology & therapy. 2015;16(3).

32. Breuleux M. Role of heregulin in human cancer. Cellular and molecular life sciences : CMLS. 2007;64(18).

33. Datta S, Dudek H, Tao X, Masters S, Fu H, Gotoh Y, et al. Akt phosphorylation of BAD couples survival signals to the cell-intrinsic death machinery. Cell. 1997;91(2).

34. Cheng J, Jiang X, Fraser M, Li M, Dan H, Sun M, et al. Role of X-linked inhibitor of apoptosis protein in chemoresistance in ovarian cancer: possible involvement of the phosphoinositide-3 kinase/Akt pathway. Drug resistance updates : reviews and commentaries in antimicrobial and anticancer chemotherapy. 2002;5(3-4).

35. Anandharaj A, Cinghu S, Park W. Rapamycin-mediated mTOR inhibition attenuates survivin and sensitizes glioblastoma cells to radiation therapy. Acta biochimica et biophysica Sinica. 2011;43(4).

36. Yang L, Y L, Shen E, Cao F, Li L, Li X, et al. NRG1-dependent activation of HER3 induces primary resistance to trastuzumab in HER2-overexpressing breast cancer cells. International journal of oncology. 2017;51(5).

37. Shibuya M, Komi E, Wang R, Kato T, Watanabe Y, Sakai M, et al. Measurement and comparison of serum neuregulin 1 immunoreactivity in control subjects and patients with schizophrenia: an influence of its genetic polymorphism. Journal of neural transmission (Vienna, Austria : 1996). 2010;117(7).

38. Hardwick JM, Soane L. Multiple functions of BCL-2 family proteins. Cold Spring Harb Perspect Biol. 2013;5(2).

39. Singh R, Letai A, Sarosiek K. Regulation of apoptosis in health and disease: the balancing act of BCL-2 family proteins. Nature reviews Molecular cell biology. 2019;20(3).

40. Zimmer AS, Van Swearingen AED, Anders CK. HER2-positive breast cancer brain metastasis: A new and exciting landscape. Cancer Rep (Hoboken). 2022;5(4):e1274.

41. Kuksis M, Gao Y, Tran W, Hoey C, Kiss A, Komorowski AS, et al. The incidence of brain metastases among patients with metastatic breast cancer: a systematic review and meta-analysis. Neuro Oncol. 2021;23(6):894–904.

42. Lin N, Winer E. Brain metastases: the HER2 paradigm. Clinical cancer research : an official journal of the American Association for Cancer Research. 2007;13(6).

43. Hurvitz SA, O’Shaughnessy J, Mason G, Yardley DA, Jahanzeb M, Brufsky A, et al. Central Nervous System Metastasis in Patients with HER2-Positive Metastatic Breast Cancer: Patient Characteristics, Treatment, and Survival from SystHERs. Clin Cancer Res. 2019;25(8):2433–41.

44. Kuksis M, Gao Y, Tran W, Hoey C, Kiss A, Komorowski A, et al. The incidence of brain metastases among patients with metastatic breast cancer: a systematic review and meta-analysis. Neuro-oncology. 2021;23(6).

45. O’Sullivan CC, Davarpanah NN, Abraham J, Bates SE. Current challenges in the management of breast cancer brain metastases. Semin Oncol. 2017;44(2):85–100.

46. Dai X, Cheng H, Bai Z, Li J. Breast Cancer Cell Line Classification and Its Relevance with Breast Tumor Subtyping. Journal of Cancer. 2017;8(16).

47. L M, M V. Animal models of brain metastasis. Neuro-oncology advances. 2021;3(Suppl 5).

48. Buonanno A, Fischbach G. Neuregulin and ErbB receptor signaling pathways in the nervous system. Current opinion in neurobiology. 2001;11(3).

49. Mei L, Nave K. Neuregulin-ERBB signaling in the nervous system and neuropsychiatric diseases. Neuron. 2014;83(1).

50. Fernald K, Kurokawa M. Evading apoptosis in cancer. Trends in cell biology. 2013;23(12).

51. Mann J, Githaka JM, Buckland TW, Yang N, Montpetit R, Patel N, et al. Non-canonical BAD activity regulates breast cancer cell and tumor growth via 14-3-3 binding and mitochondrial metabolism. Oncogene. 2019;38(18):3325–39.

52. Kim TM, Lee KW, Oh DY, Lee JS, Im SA, Kim DW, et al. Phase 1 Studies of Poziotinib, an Irreversible Pan-HER Tyrosine Kinase Inhibitor in Patients with Advanced Solid Tumors. Cancer Res Treat. 2018;50(3):835–42.

53. Elamin YY, Robichaux JP, Carter BW, Altan M, Gibbons DL, Fossella FV, et al. Poziotinib for Patients With HER2 Exon 20 Mutant Non-Small-Cell Lung Cancer: Results From a Phase II Trial. J Clin Oncol. 2022;40(7):702–9.

54. Le X, Cornelissen R, Garassino M, Clarke JM, Tchekmedyian N, Goldman JW, et al. Poziotinib in Non-Small-Cell Lung Cancer Harboring HER2 Exon 20 Insertion Mutations After Prior Therapies: ZENITH20-2 Trial. J Clin Oncol. 2022;40(7):710–8.

55. Qi X, Shi Q, Xuhong J, Zhang Y, Jiang J. Pyrotinib-based therapeutic approaches for HER2 positive breast cancer: the time is now. Breast Cancer Res. 2023;25(1):113.

56. Gupta M, HJ L, Barden C, Weaver D. The Blood-Brain Barrier (BBB) Score. Journal of medicinal chemistry. 2019;62(21).

57. F M, J C, PW A. Predictors for Blood-Brain Barrier Permeability with resampling strategies based on B3DB database. 2019.

58. SumulationsPlus. ADMET Predictor®.

59. Franken N, Rodermond H, Stap J, Haveman J, van Bree C. Clonogenic assay of cells in vitro. Nature protocols. 2006;1(5).

60. Livak K, Schmittgen T. Analysis of relative gene expression data using real-time quantitative PCR and the 2(-Delta Delta C(T)) Method. Methods (San Diego, Calif). 2001;25(4).

61. ACDBio. RNAscope Sample Preparation and Pretreatment Guide for FFPE Tissue 2013 [

62. ACDBio. RNAscope 2.5 HD Duplex Detection Kit (Chromogenic) 2019 [

63. Baumann BC, Dorsey JF, Benci JL, Joh DY, Kao GD. Stereotactic intracranial implantation and in vivo bioluminescent imaging of tumor xenografts in a mouse model system of glioblastoma multiforme. J Vis Exp. 2012(67).

